## Supplementary Information for "Molecular mechanism of exchange coupling in CLC chloride/proton antiporters"

### Supplementary Table 1. Cryo-EM data collection, refinement and validation statistics

| | CLC-ec1 pH 7.5 | CLC-ec1 pH 4 | CLC-ec1 pH 3 | CLC-ec1 K131A pH 7.5 |
|---|---|---|---|---|
| **Data collection and processing** | | | | |
| Magnification | 96,000 | 105,000 | 165,000 | 130,000 |
| Voltage (kV) | 300 | 300 | 300 | 300 |
| Electron exposure (e–/Å$^2$) | 80 | 60 | 60 | 60 |
| Defocus range (μm) | -0.8 to -2 | -0.8 to -2 | -0.8 to -2 | -0.8 to -2 |
| Pixel size (Å) | 0.82 | 0.8642 | 0.741 | 0.68 |
| Symmetry imposed | C2 | C2 | C2 | C2 |
| Initial particle images (no.) | 971,482 | 871,809 | 1,556,961 | 591,019 |
| Final particle images (no.) | 72,103 | 261,507 | 66,046 | 307,722 |
| Map resolution (Å) | 3.2 | 3.2 | 3.1 | 3.3 |
| FSC threshold | 0.143 | 0.143 | 0.143 | 0.143 |
| Map resolution range (Å) | 2 to 7.9 | 2.1 to 9.5 | 1.8 to 7.2 | 2.1 to 7.8 |
| **Refinement** | | | | |
| Initial model used (PDB code) | 1OTS | 1OTS | 6V2J | 1OTS |
| Model resolution (Å) | 3.1 | 3.1 | 3.1 | 3.3 |
| FSC threshold | 0.143 | 0.143 | 0.143 | 0.143 |
| Map sharpening *B* factor (Å$^2$) | -147.2 | -115.9 | -116.9 | -136.9 |
| Model composition | | | | |
| Non-hydrogen atoms | 6668 | 6670 | 6486 | 6660 |
| Protein residues | 888 | 888 | 870 | 888 |
| Ligands | 2 | 2 | 4 | 2 |
| *B* factors (Å$^2$) | | | | |
| Protein | 50.12 | 49.39 | 59.92 | 43.67 |
| Ligand | 70.43 | 59.92 | 86.14 | 88.32 |
| R.m.s. deviations | | | | |
| Bond lengths (Å) | 0.003 | 0.002 | 0.002 | 0.003 |
| Bond angles (°) | 0.511 | 0.535 | 0.529 | 0.541 |
| **Validation** | | | | |
| MolProbity score | 1.01 | 0.97 | 1.22 | 1.07 |
| Clashscore | 2.35 | 1.98 | 3.17 | 1.98 |
| Poor rotamers (%) | 0.30 | 0.60 | 0.62 | 0.60 |
| Ramachandran plot | | | | |
| Favored (%) | 98.42 | 97.96 | 97.46 | 97.51 |
| Allowed (%) | 1.58 | 2.04 | 2.54 | 2.49 |
| Disallowed (%) | 0.00 | 0.00 | 0.00 | 0.00 |

**Supplementary Table 2. Simulations Setup Details**

| | |
|---|---|
| Box Dimensions* | 160 Å x120 Å x90 Å |
| Salt Concentration | 150 mM sodium ions, 150 mM chloride ions |
| Total Atoms* | 123,000 |
| Total Waters* | 24,000 |
| Total Lipids* | 280 |
| Lipid Type | palmitoyl-oleoyl-phosphatidylcholine (POPC) |

*Approximate values are listed for box dimensions, total atoms, total waters, and total lipids, as precise values vary between simulation conditions.

### Supplementary Table 3. Reliability and reproducibility checklist for MD simulations

| 1. Convergence of simulations and analysis | Yes | N/A | |
|---|---|---|---|
| 1a. Is an evaluation presented in the text to show that the property being measured has equilibrated in the simulations (*e.g.* time-course analysis)? | ☐ | | We present time-course analysis for key metrics (Figure 4c, Extended Data Figure 9) and show that behavior is consistent across independent simulations. |
| 1b. Then, is it described in the text how simulations are split into equilibration and production runs and how much data were analyzed from production runs? | ☒ | | Equilibration and production simulation procedures are described in the "Molecular dynamics simulation and analysis protocols" portion of the Methods section. |
| 1c. Are there at least 3 simulations per simulation condition with statistical analysis? | ☒ | | Ten simulations were performed per condition, as described in the Methods section. |
| 1d. Is evidence provided in the text that the simulation results presented are independent of initial configuration? | ☒ | | Each independent simulation was subjected to its own equilibration steps and initialized with different velocities. Thus, each simulation has a distinct initial configuration. |
| **2. Connection to experiments** | | | |
| 2a. Are calculations provided that can connect to experiments (*e.g.* loss or gain in function from mutagenesis, binding assays, NMR chemical shifts, J-couplings, SAXS curves, interaction distances or FRET distances, structure factors, diffusion coefficients, bulk modulus and other mechanical properties, *etc.*)? | ☒ | | The Discussion sections of our manuscript titled "Conformational dynamics revealed by molecular dynamics simulations" and "The mechanism of reversible $Cl^-/H^+$ exchange" explain how simulation results connect to experimental ion exchange and mutagenesis data. |
| **3. Method choice** | | | |
| 3a. Do simulations contain membranes, membrane proteins, intrinsically disordered proteins, glycans, nucleic acids, polymers, or cryptic ligand binding? | ☒ | ☐ | Simulations contain receptors embedded in lipid membranes as described in the Methods section "System setup for MD simulations." |
| 3b. Is it described in the text whether the accuracy of the chosen model(s) is sufficient to address the question(s) under investigation (e.g., all-atom vs. coarse-grained models, fixed charge vs. polarizable force fields, implicit vs. explicit solvent or membrane, specific force field and water model, etc.? | ☐ | | We explain the justification for and limitations of our simulation setup within the Discussion section titled "The mechanism of reversible $Cl^-/H^+$ exchange." |
| 3c. Is the timescale of the event(s) under investigation beyond the brute-force MD simulation timescale in this study that enhanced sampling methods are needed? | ☐ | ☒ | |
| If **NO**, is the evidence provided in the text? | ☒ | | Time-course analyses (Figure 4c, Supplementary Figure 9) show that simulations have converged for the metrics of interest. |
| **4. Code and reproducibility** | | | |
| 4a. Is a table provided describing the system setup, such as simulation box dimensions, total number of atoms, total number of water molecules, salt concentration, lipid composition (number of molecules and type)? | ☒ | | This information is listed in Supplementary Table 2. |
| 4b. Are other parameters for the system setup described in the text, such as protonation state, type of structural restraints if applied, nonbonded cutoff, thermostat and barostat, etc.? | ☒ | | All parameters are described within the Methods section (specifically, within "System setup for molecular dynamics simulations" and "Molecular dynamics simulation and analysis protocols"). |
| 4c. Is it described in the text what simulation and analysis software and which versions are used? | ☒ | | All software packages and versions used are described within the Methods section (specifically, within "System setup for molecular dynamics simulations" and "Molecular dynamics simulation and analysis protocols"). |
| 4d. Are initial coordinate and simulation input files and a coordinate file of the final output provided as supplementary files or in a public repository? | ☒ | | The initial coordinate file, simulation input files, and trajectories are available on Zenodo (10.5281/zenodo.17808101) |
| 4e. Is there custom code or custom force field parameters? | ☐ | ☒ | Not applicable. |

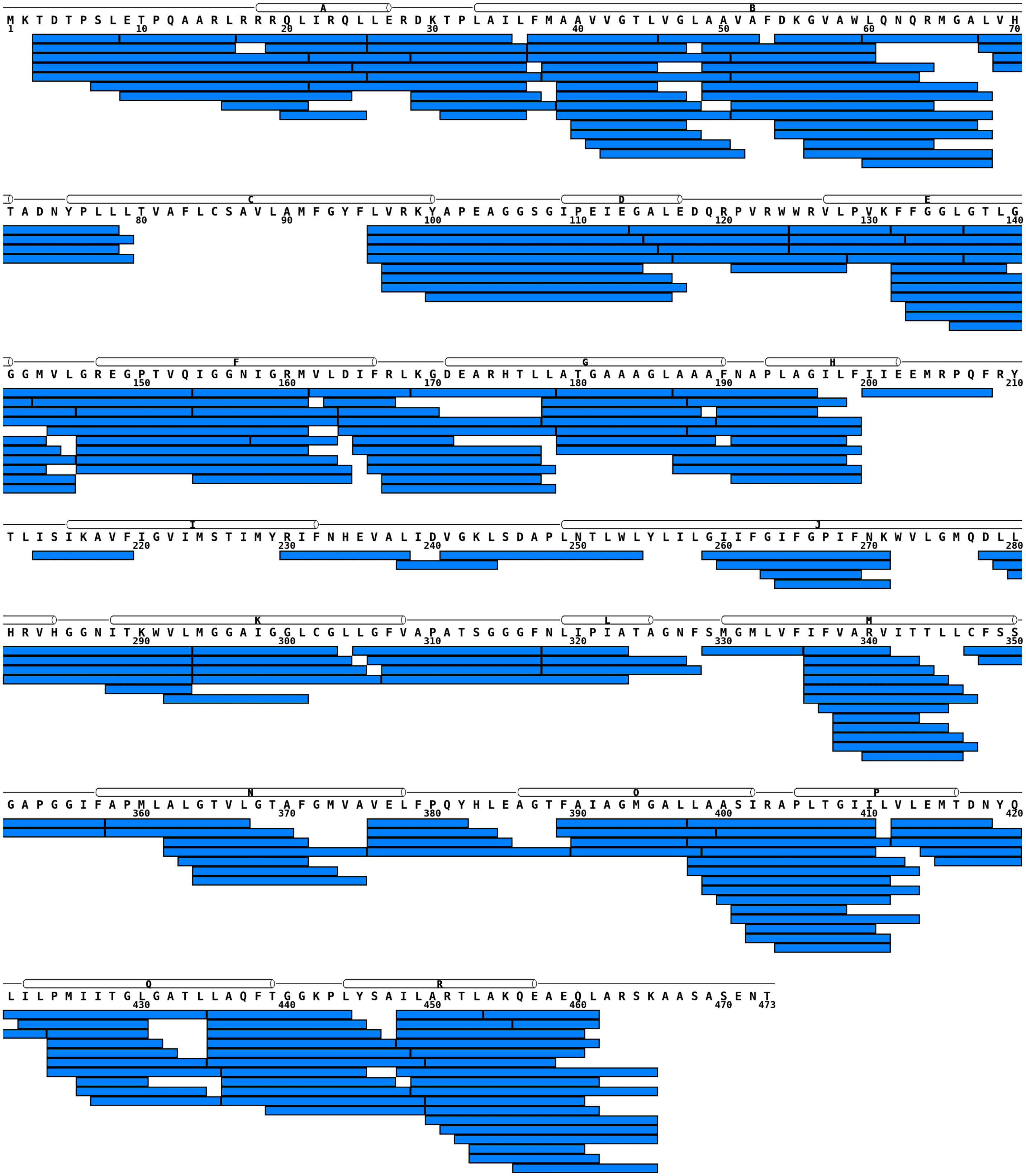

**Supplementary Figure 1. Sequence coverage map of CLC-ec1 by peptides providing HDX data.** Online proteolysis of CLC-ec1 using co-immobilized Nepenthesin-2/Pepsin column (bed volume 70 µl). Digestion conditions: 19°C, 0.4% formic acid, flow rate of 200 µL min$^{-1}$. Each peptide is represented by a blue bar. Secondary structure elements are shown above the sequence. Almost complete (89%) sequence coverage was obtained through 237 peptides with average peptide length of 11.4 amino acids and average redundancy of 6.4.

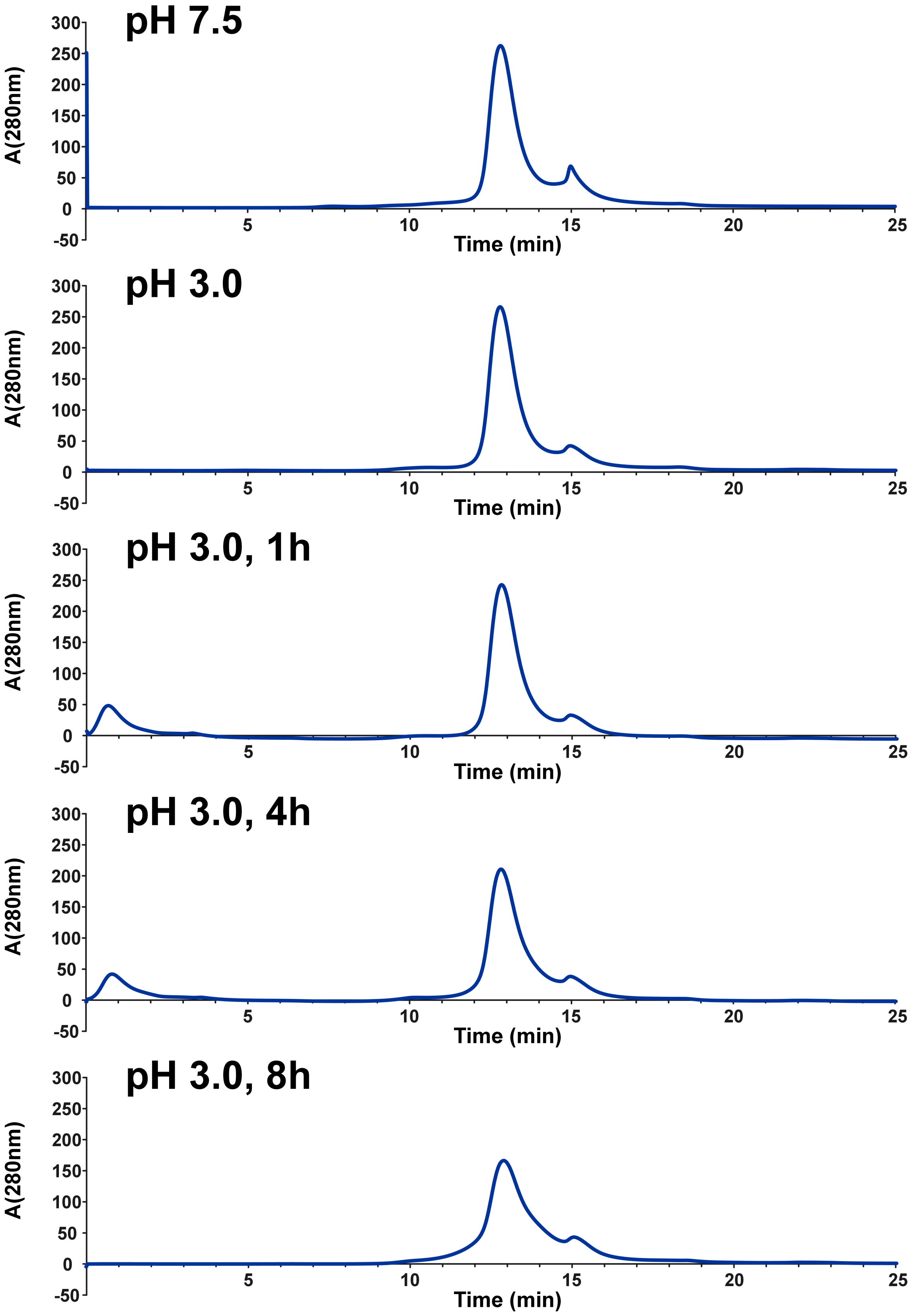

**Supplementary Figure 2: CLC-ec1 stability at pH 3.0.** Size exclusion chromatography of CLC-ec1 using EnrichTM SEC 650 10x300 24 mL (Biorad), flow rate 1 mL/min. The top panel shows CLC-ec1 prepared at pH 7.5. The lower panels show CLC-ec1 assessed using size exclusion chromatography immediately after preparation at pH 3.0 and at several time points, demonstrating stability over an 8-hour period.

**Supplementary Figure 3A: Complete set of deuterium uptake curves from the HDX-MS experiment – Part A.**

Deuterium uptake plots for all CLC-ec1 peptides providing HDX data. Five experimental conditions were followed: pH 3.0 (black), pH 3.5 (grey), pH 4.0 (blue), pH 4.5 (cyan), and pH 6.5 (orange). No correction for the intrinsic pH-dependent exchange rate differences was applied. Samples were collected at 20s, 63s, 200s, 633s, 2000s, 6325s, 20000s, and 63000s. The 20s time point contains a technical triplicate combined with an additional triplicate representing protein stability control. For this control, CLC-ec1 was incubated in $H_2O$ buffer of the respective pH for 63000s (equivalent to the longest time point) and subsequently subjected to a 20s $D_2O$ labeling pulse. Mean values with corresponding standard deviations are shown in the plots. A fully deuterated control was used to correct for back-exchange and compensate for deuterium loss during analysis. Peptide limits and charge states are indicated above each plot.

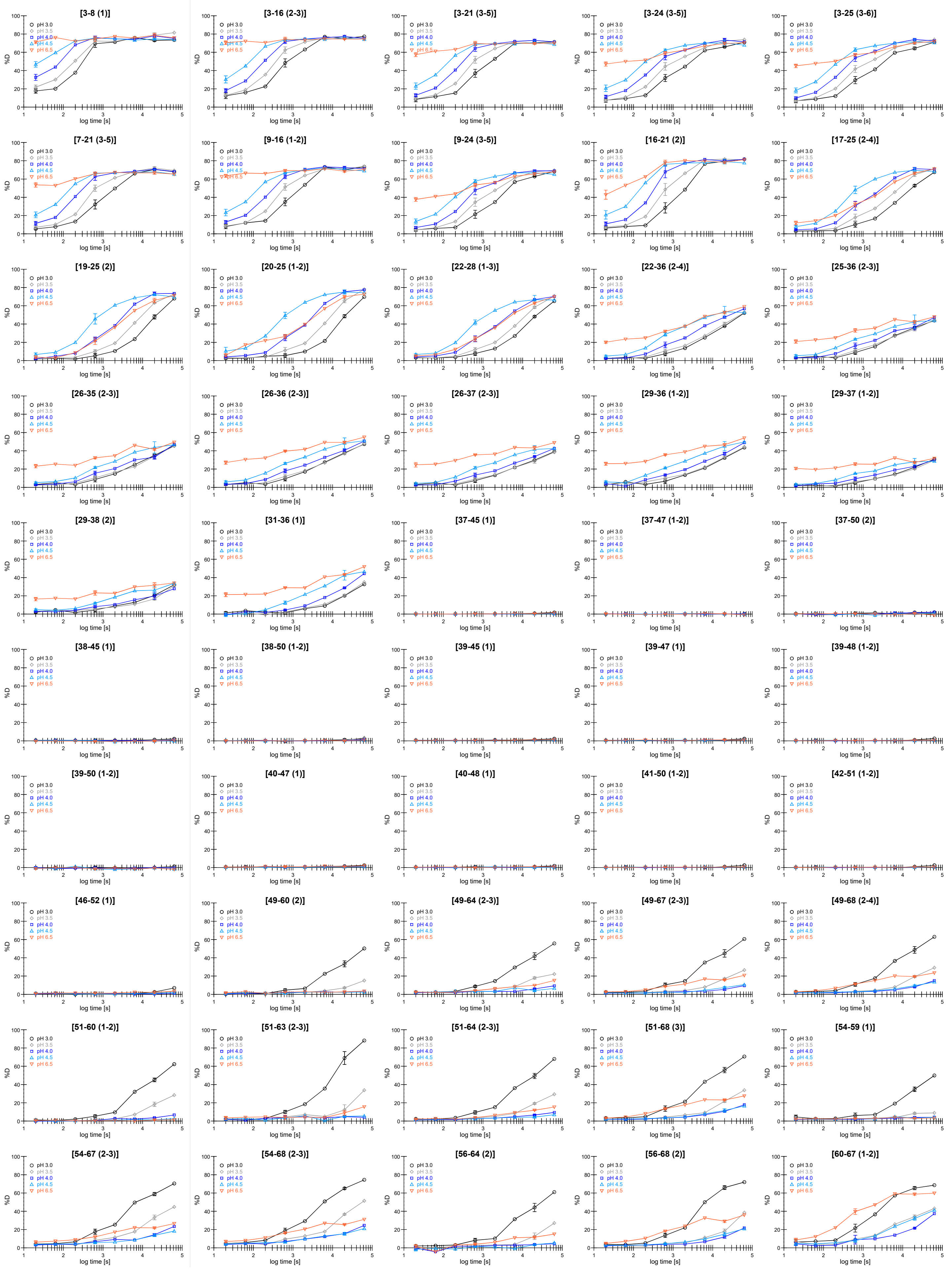
[3-8 (1)]
[3-16 (2-3)]
[3-21 (3-5)]
[3-24 (3-5)]
[3-25 (3-6)]
[7-21 (3-5)]
[9-16 (1-2)]
[9-24 (3-5)]
[16-21 (2)]
[17-25 (2-4)]
[19-25 (2)]
[20-25 (1-2)]
[22-28 (1-3)]
[22-36 (2-4)]
[25-36 (2-3)]
[26-35 (2-3)]
[26-36 (2-3)]
[26-37 (2-3)]
[29-36 (1-2)]
[29-37 (1-2)]
[29-38 (2)]
[31-36 (1)]
[37-45 (1)]
[37-47 (1-2)]
[37-50 (2)]
[38-45 (1)]
[38-50 (1-2)]
[39-45 (1)]
[39-47 (1)]
[39-48 (1-2)]
[39-50 (1-2)]
[40-47 (1)]
[40-48 (1)]
[41-50 (1-2)]
[42-51 (1-2)]
[46-52 (1)]
[49-60 (2)]
[49-64 (2-3)]
[49-67 (2-3)]
[49-68 (2-4)]
[51-60 (1-2)]
[51-63 (2-3)]
[51-64 (2-3)]
[51-68 (3)]
[54-59 (1)]
[54-67 (2-3)]
[54-68 (2-3)]
[56-64 (2)]
[56-68 (2)]
[60-67 (1-2)]
pH 3.0
pH 3.5
pH 4.0
pH 4.5
pH 6.5
%D
log time [s]

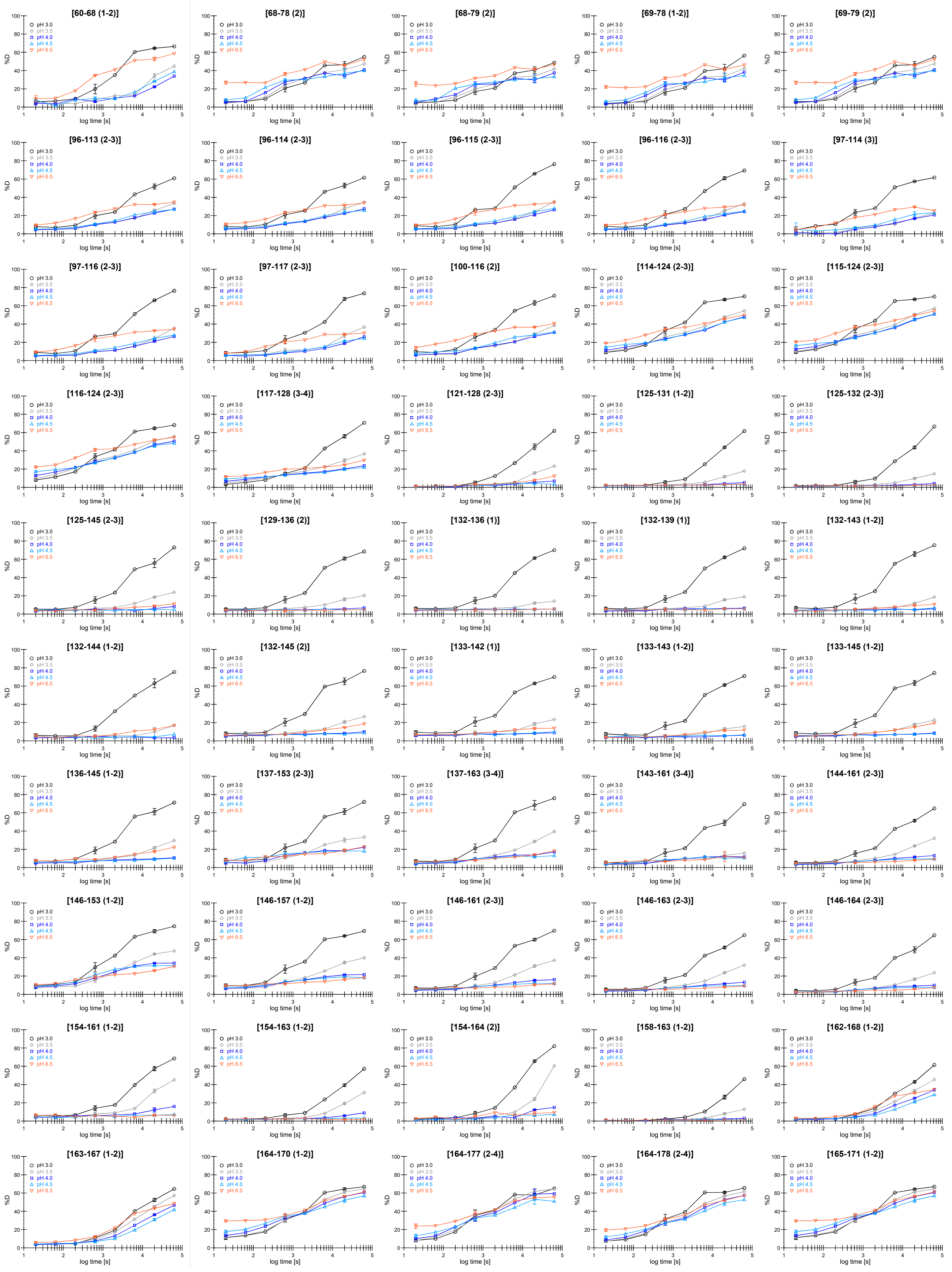
[60-68 (1-2)]
[68-78 (2)]
[68-79 (2)]
[69-78 (1-2)]
[69-79 (2)]
[96-113 (2-3)]
[96-114 (2-3)]
[96-115 (2-3)]
[96-116 (2-3)]
[97-114 (3)]
[97-116 (2-3)]
[97-117 (2-3)]
[100-116 (2)]
[114-124 (2-3)]
[115-124 (2-3)]
[116-124 (2-3)]
[117-128 (3-4)]
[121-128 (2-3)]
[125-131 (1-2)]
[125-132 (2-3)]
[125-145 (2-3)]
[129-136 (2)]
[132-136 (1)]
[132-139 (1)]
[132-143 (1-2)]
[132-144 (1-2)]
[132-145 (2)]
[133-142 (1)]
[133-143 (1-2)]
[133-145 (1-2)]
[136-145 (1-2)]
[137-153 (2-3)]
[137-163 (3-4)]
[143-161 (3-4)]
[144-161 (2-3)]
[146-153 (1-2)]
[146-157 (1-2)]
[146-161 (2-3)]
[146-163 (2-3)]
[146-164 (2-3)]
[154-161 (1-2)]
[154-163 (1-2)]
[154-164 (2)]
[158-163 (1-2)]
[162-168 (1-2)]
[163-167 (1-2)]
[164-170 (1-2)]
[164-177 (2-4)]
[164-178 (2-4)]
[165-171 (1-2)]
pH 3.0
pH 3.5
pH 4.0
pH 4.5
pH 6.5
%D
log time [s]

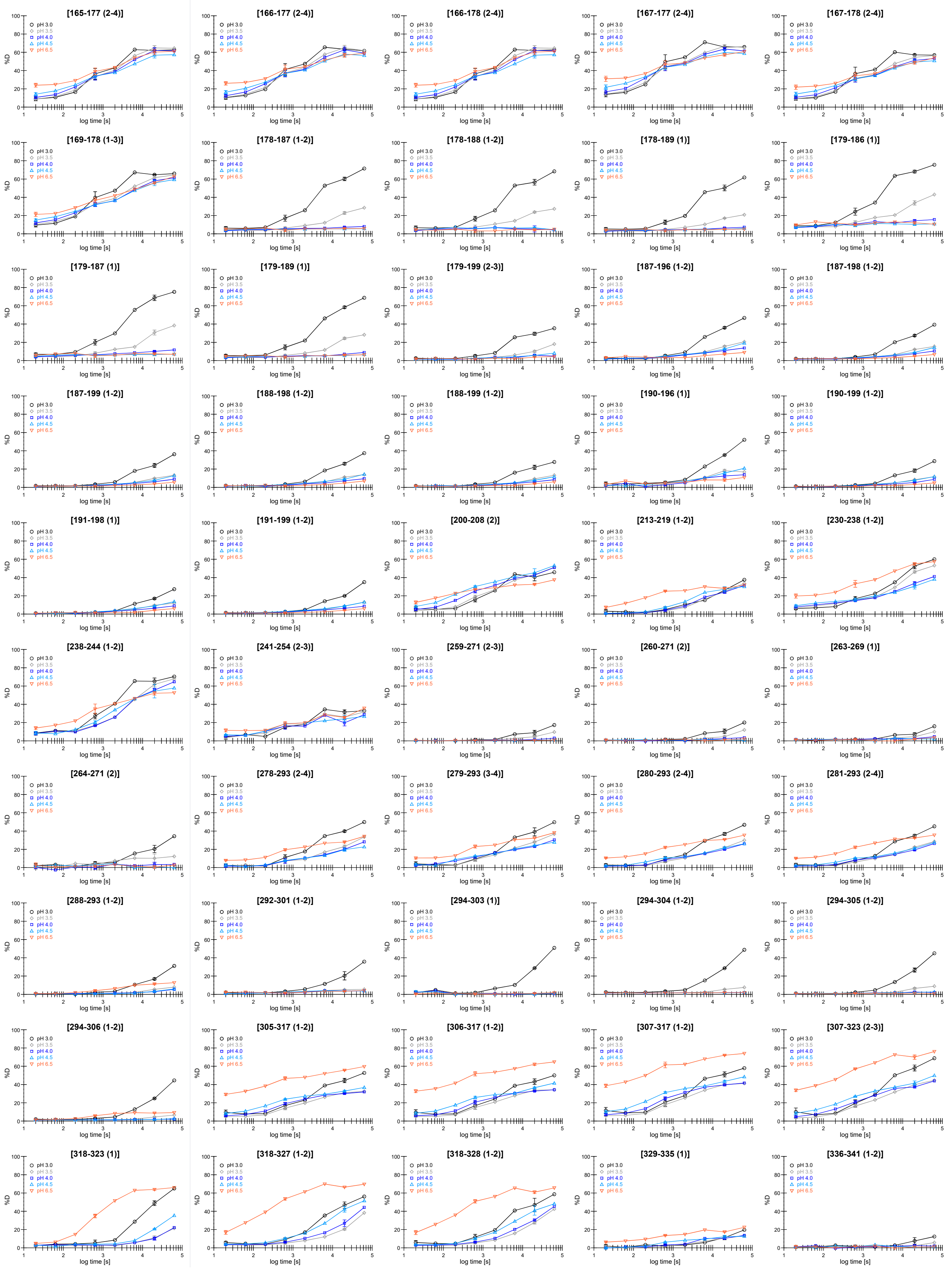
[165-177 (2-4)]
[166-177 (2-4)]
[166-178 (2-4)]
[167-177 (2-4)]
[167-178 (2-4)]
[169-178 (1-3)]
[178-187 (1-2)]
[178-188 (1-2)]
[178-189 (1)]
[179-186 (1)]
[179-187 (1)]
[179-189 (1)]
[179-199 (2-3)]
[187-196 (1-2)]
[187-198 (1-2)]
[187-199 (1-2)]
[188-198 (1-2)]
[188-199 (1-2)]
[190-196 (1)]
[190-199 (1-2)]
[191-198 (1)]
[191-199 (1-2)]
[200-208 (2)]
[213-219 (1-2)]
[230-238 (1-2)]
[238-244 (1-2)]
[241-254 (2-3)]
[259-271 (2-3)]
[260-271 (2)]
[263-269 (1)]
[264-271 (2)]
[278-293 (2-4)]
[279-293 (3-4)]
[280-293 (2-4)]
[281-293 (2-4)]
[288-293 (1-2)]
[292-301 (1-2)]
[294-303 (1)]
[294-304 (1-2)]
[294-305 (1-2)]
[294-306 (1-2)]
[305-317 (1-2)]
[306-317 (1-2)]
[307-317 (1-2)]
[307-323 (2-3)]
[318-323 (1)]
[318-327 (1-2)]
[318-328 (1-2)]
[329-335 (1)]
[336-341 (1-2)]
pH 3.0
pH 3.5
pH 4.0
pH 4.5
pH 6.5
%D
log time [s]

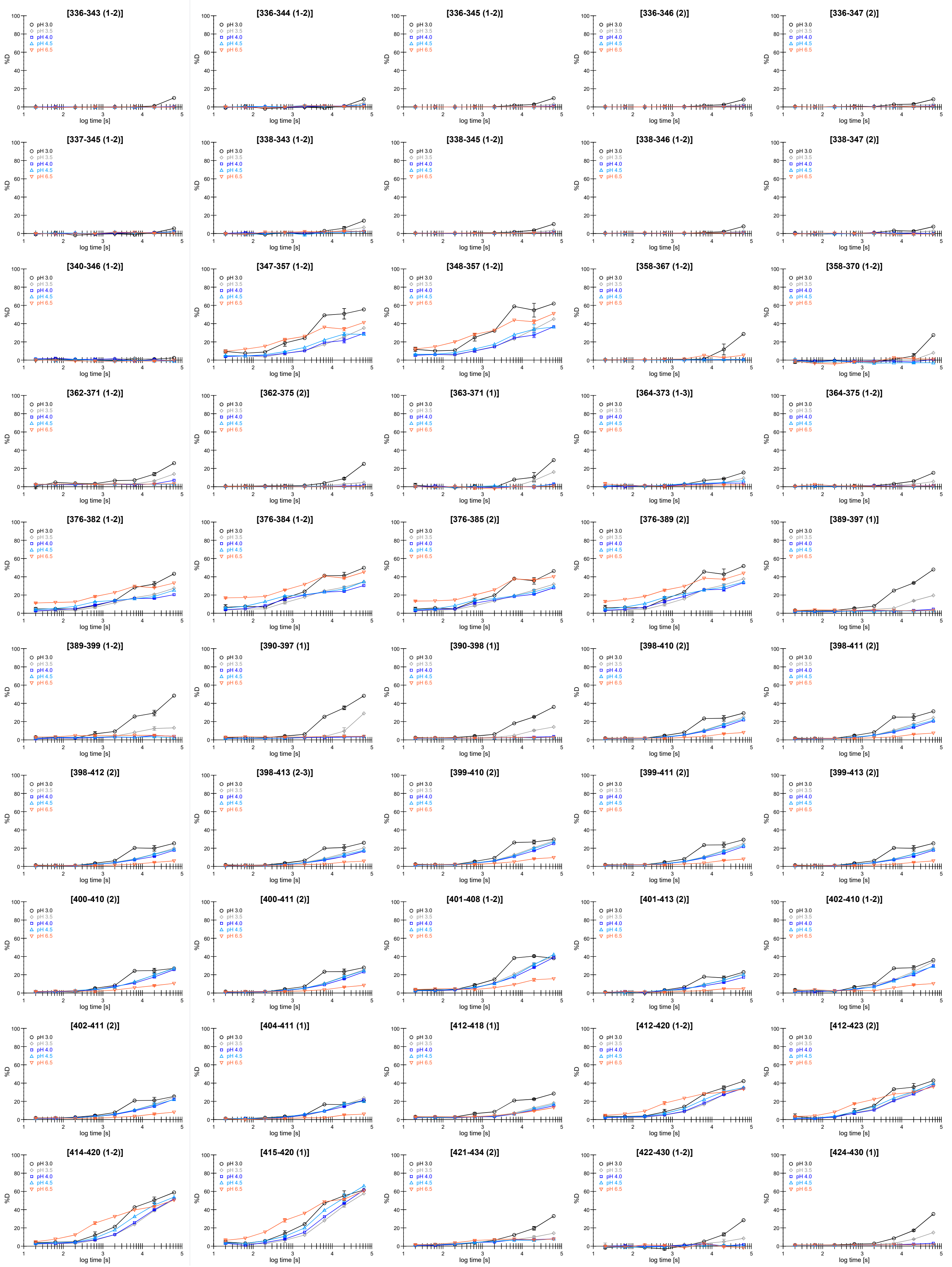
[336-343 (1-2)]
[336-344 (1-2)]
[336-345 (1-2)]
[336-346 (2)]
[336-347 (2)]
[337-345 (1-2)]
[338-343 (1-2)]
[338-345 (1-2)]
[338-346 (1-2)]
[338-347 (2)]
[340-346 (1-2)]
[347-357 (1-2)]
[348-357 (1-2)]
[358-367 (1-2)]
[358-370 (1-2)]
[362-371 (1-2)]
[362-375 (2)]
[363-371 (1)]
[364-373 (1-3)]
[364-375 (1-2)]
[376-382 (1-2)]
[376-384 (1-2)]
[376-385 (2)]
[376-389 (2)]
[389-397 (1)]
[389-399 (1-2)]
[390-397 (1)]
[390-398 (1)]
[398-410 (2)]
[398-411 (2)]
[398-412 (2)]
[398-413 (2-3)]
[399-410 (2)]
[399-411 (2)]
[399-413 (2)]
[400-410 (2)]
[400-411 (2)]
[401-408 (1-2)]
[401-413 (2)]
[402-410 (1-2)]
[402-411 (2)]
[404-411 (1)]
[412-418 (1)]
[412-420 (1-2)]
[412-423 (2)]
[414-420 (1-2)]
[415-420 (1)]
[421-434 (2)]
[422-430 (1-2)]
[424-430 (1)]
pH 3.0
pH 3.5
pH 4.0
pH 4.5
pH 6.5
%D
log time [s]

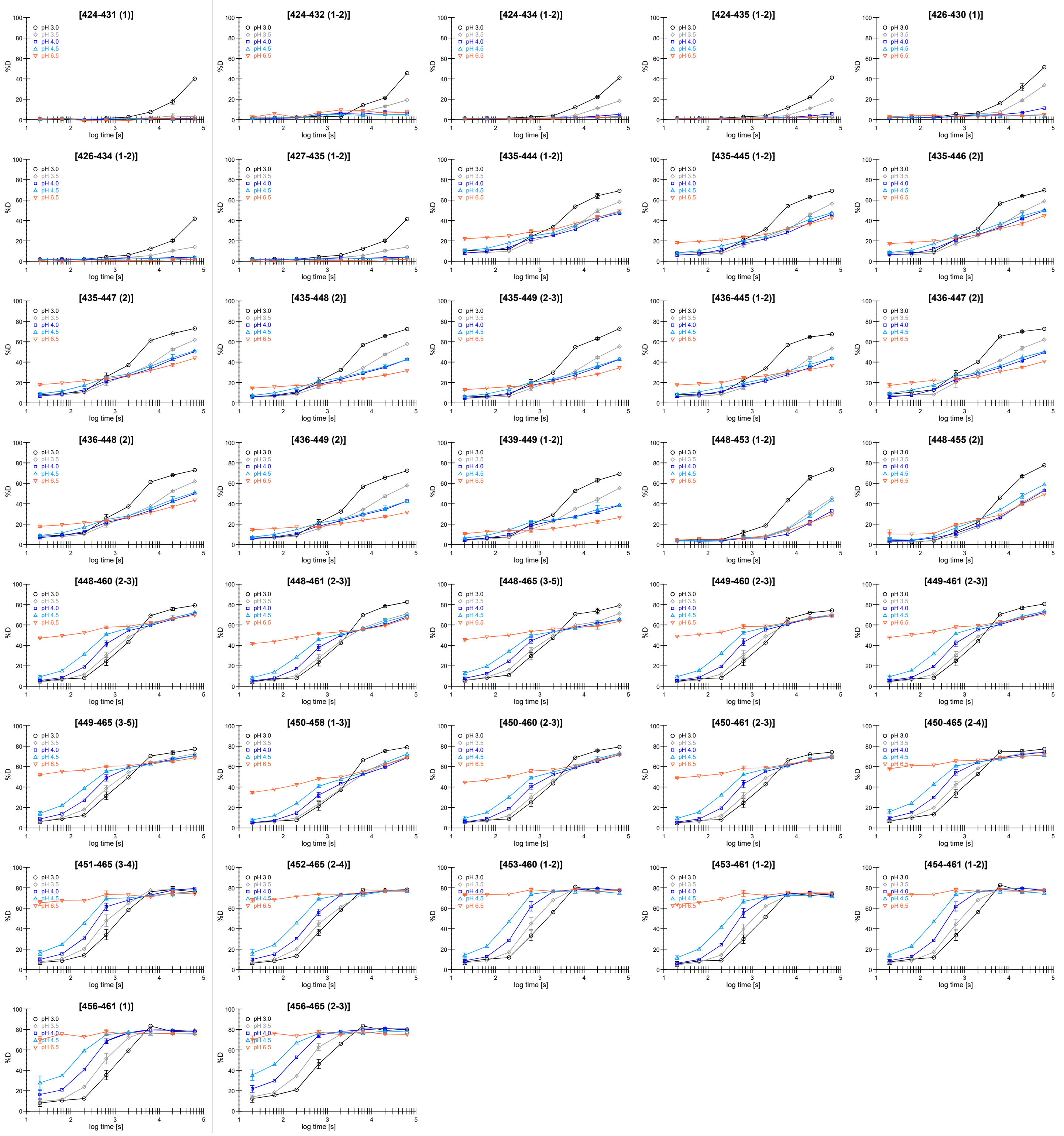
[424-431 (1)]
[424-432 (1-2)]
[424-434 (1-2)]
[424-435 (1-2)]
[426-430 (1)]
[426-434 (1-2)]
[427-435 (1-2)]
[435-444 (1-2)]
[435-445 (1-2)]
[435-446 (2)]
[435-447 (2)]
[435-448 (2)]
[435-449 (2-3)]
[436-445 (1-2)]
[436-447 (2)]
[436-448 (2)]
[436-449 (2)]
[439-449 (1-2)]
[448-453 (1-2)]
[448-455 (2)]
[448-460 (2-3)]
[448-461 (2-3)]
[448-465 (3-5)]
[449-460 (2-3)]
[449-461 (2-3)]
[449-465 (3-5)]
[450-458 (1-3)]
[450-460 (2-3)]
[450-461 (2-3)]
[450-465 (2-4)]
[451-465 (3-4)]
[452-465 (2-4)]
[453-460 (1-2)]
[453-461 (1-2)]
[454-461 (1-2)]
[456-461 (1)]
[456-465 (2-3)]
pH 3.0
pH 3.5
pH 4.0
pH 4.5
pH 6.5
%D
log time [s]

**Supplementary Figure 3B: Complete set of deuterium uptake curves from the HDX-MS experiment – Part B**

Deuterium uptake plots for all CLC-ec1 peptides providing HDX data. Five experimental conditions were followed: pH 3.0 (black), pH 3.5 (grey), pH 4.0 (blue), pH 4.5 (cyan), and pH 6.5 (orange). These plots are normalized to the exchange rate at pH 3.0. The commonly used linear approximation (10-fold change per pH unit) is valid only in the upper range (e.g., pH 6.5 vs. 4.5) but deviates significantly at lower pH values. Therefore, theoretical intrinsic exchange rates were calculated using the SPHERE server (https://spin.niddk.nih.gov/bax-apps/nmrserver/sphere/index.html) based on previously established reference rates [1, 2]. Calculations were performed with the following settings: temperature 21 °C, pH meter readings 2.6, 3.1, 3.6, 4.1, and 6.1 (for pD 3.0, 3.5, 4.0, 4.5 and 6.5, respectively); reference data – alanine in oligopeptide. Exchange rates ($k_{ex}$) were obtained for each peptide bond (excluding those containing prolines), and the ratio between pH 3.0 and the other pH values was determined per bond. Median values across the protein were then used to derive correction factors (1.24 for pH 3.5, 2.63 for pH 4.0, 6.77 for pH 4.5, and 532.74 for pH 6.5). These factors were applied to adjust the exchange times at each pH, thereby normalizing the kinetics to pH 3.0. Although these corrections are derived from model peptide data, they agree reasonably well with values obtained for naturally occurring peptides [3]. The correction highlights how the unstructured N- and C-terminal ends align across conditions, providing clear evidence that the observed differences in their exchange rates were solely pH-driven. In the structured regions of the protein, the correction reveals the pH-dependent conformational transitions more clearly.

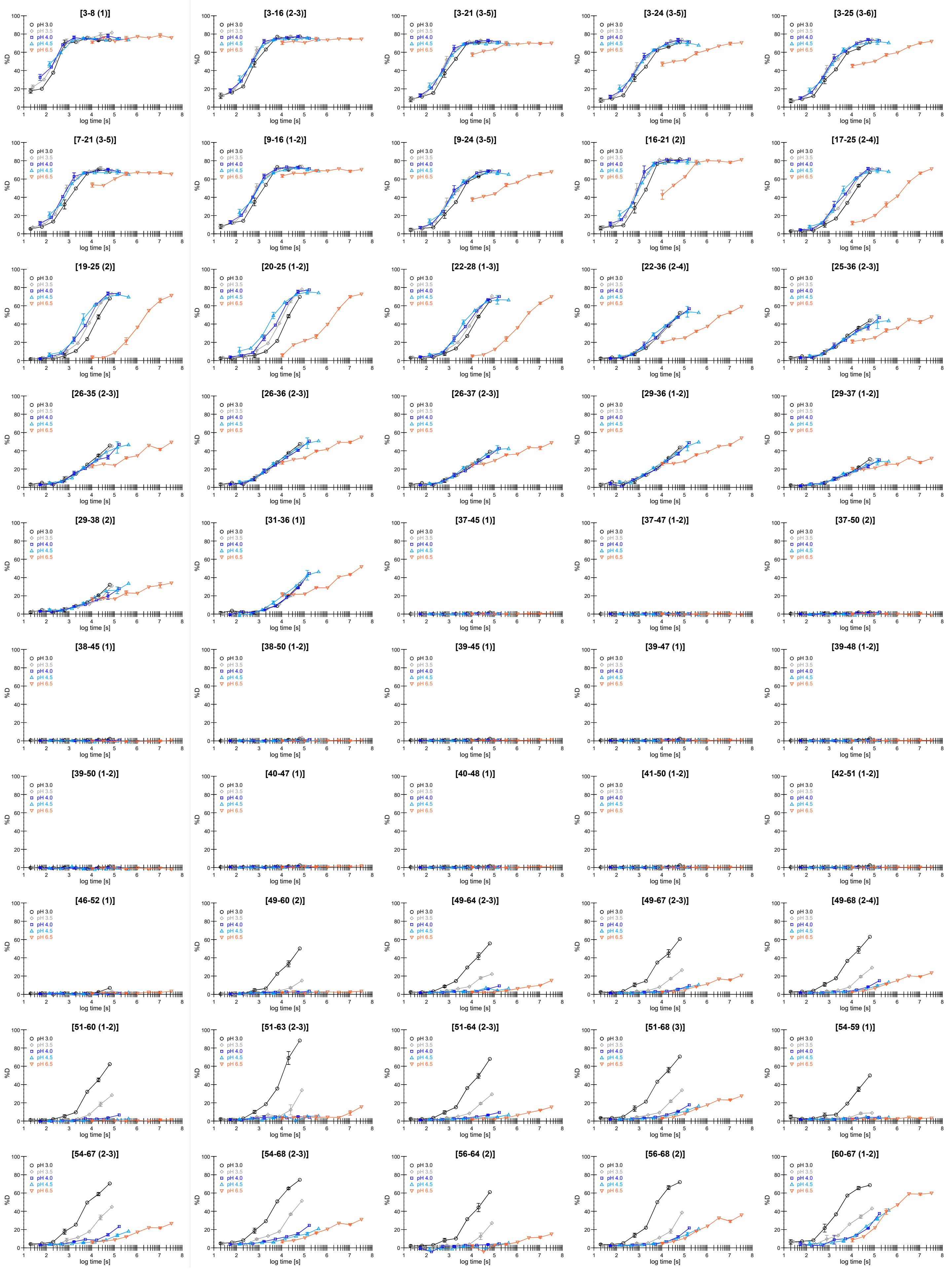
[3-8 (1)]
[3-16 (2-3)]
[3-21 (3-5)]
[3-24 (3-5)]
[3-25 (3-6)]
[7-21 (3-5)]
[9-16 (1-2)]
[9-24 (3-5)]
[16-21 (2)]
[17-25 (2-4)]
[19-25 (2)]
[20-25 (1-2)]
[22-28 (1-3)]
[22-36 (2-4)]
[25-36 (2-3)]
[26-35 (2-3)]
[26-36 (2-3)]
[26-37 (2-3)]
[29-36 (1-2)]
[29-37 (1-2)]
[29-38 (2)]
[31-36 (1)]
[37-45 (1)]
[37-47 (1-2)]
[37-50 (2)]
[38-45 (1)]
[38-50 (1-2)]
[39-45 (1)]
[39-47 (1)]
[39-48 (1-2)]
[39-50 (1-2)]
[40-47 (1)]
[40-48 (1)]
[41-50 (1-2)]
[42-51 (1-2)]
[46-52 (1)]
[49-60 (2)]
[49-64 (2-3)]
[49-67 (2-3)]
[49-68 (2-4)]
[51-60 (1-2)]
[51-63 (2-3)]
[51-64 (2-3)]
[51-68 (3)]
[54-59 (1)]
[54-67 (2-3)]
[54-68 (2-3)]
[56-64 (2)]
[56-68 (2)]
[60-67 (1-2)]
pH 3.0
pH 3.5
pH 4.0
pH 4.5
pH 6.5
%D
log time [s]

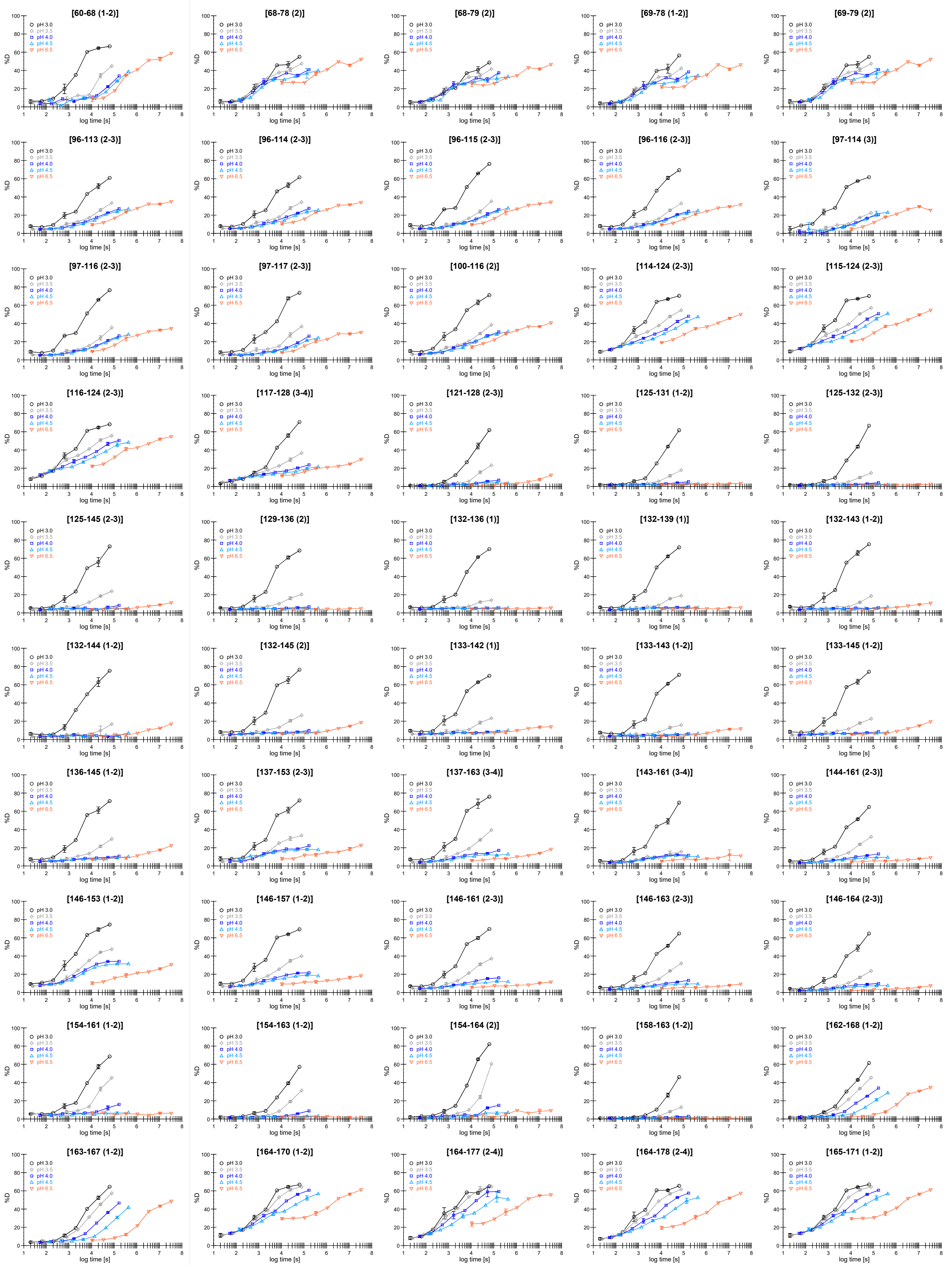
[60-68 (1-2)]
[68-78 (2)]
[68-79 (2)]
[69-78 (1-2)]
[69-79 (2)]
[96-113 (2-3)]
[96-114 (2-3)]
[96-115 (2-3)]
[96-116 (2-3)]
[97-114 (3)]
[97-116 (2-3)]
[97-117 (2-3)]
[100-116 (2)]
[114-124 (2-3)]
[115-124 (2-3)]
[116-124 (2-3)]
[117-128 (3-4)]
[121-128 (2-3)]
[125-131 (1-2)]
[125-132 (2-3)]
[125-145 (2-3)]
[129-136 (2)]
[132-136 (1)]
[132-139 (1)]
[132-143 (1-2)]
[132-144 (1-2)]
[132-145 (2)]
[133-142 (1)]
[133-143 (1-2)]
[133-145 (1-2)]
[136-145 (1-2)]
[137-153 (2-3)]
[137-163 (3-4)]
[143-161 (3-4)]
[144-161 (2-3)]
[146-153 (1-2)]
[146-157 (1-2)]
[146-161 (2-3)]
[146-163 (2-3)]
[146-164 (2-3)]
[154-161 (1-2)]
[154-163 (1-2)]
[154-164 (2)]
[158-163 (1-2)]
[162-168 (1-2)]
[163-167 (1-2)]
[164-170 (1-2)]
[164-177 (2-4)]
[164-178 (2-4)]
[165-171 (1-2)]
pH 3.0
pH 3.5
pH 4.0
pH 4.5
pH 6.5
%D
log time [s]

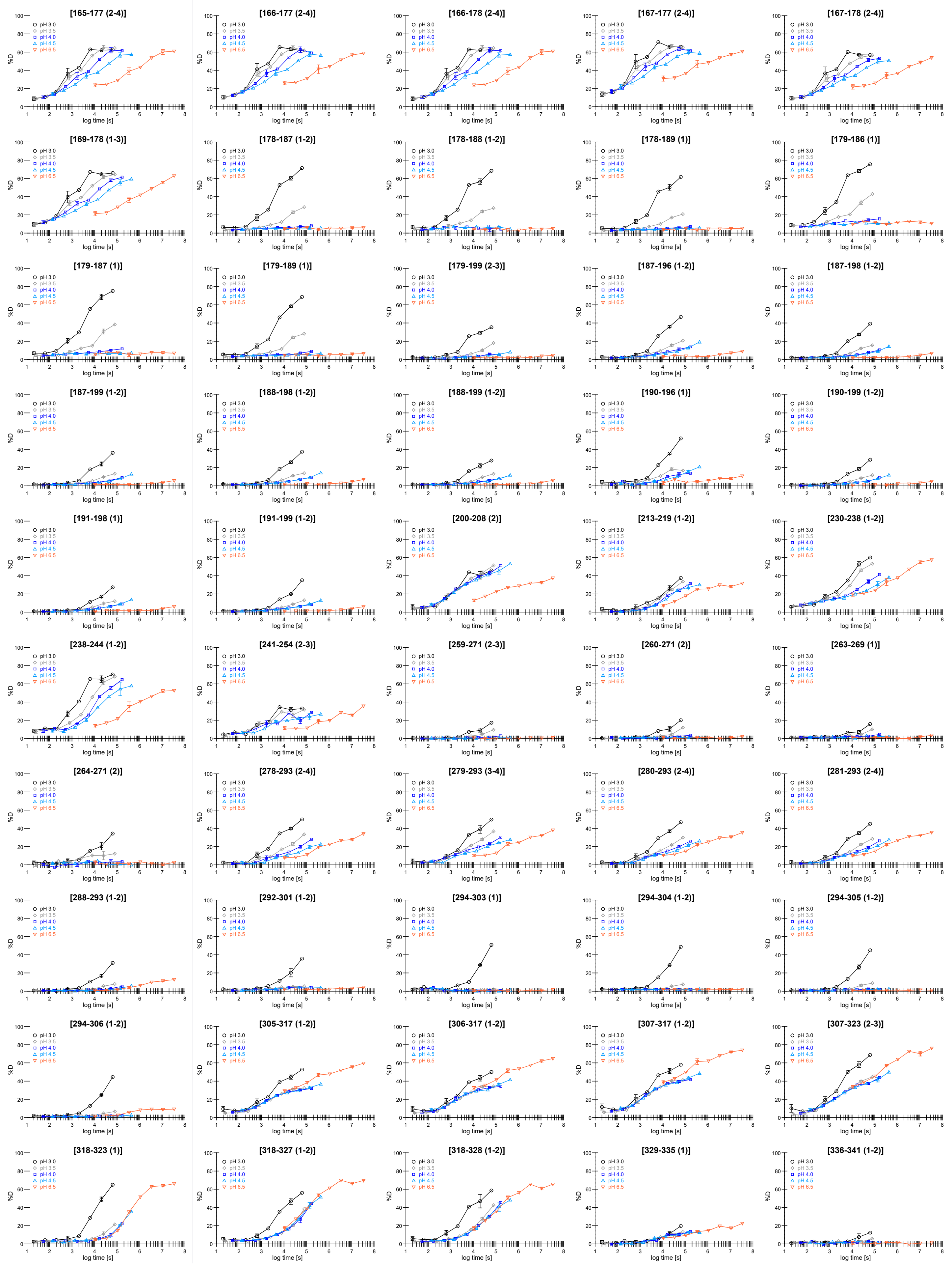
[165-177 (2-4)]
[166-177 (2-4)]
[166-178 (2-4)]
[167-177 (2-4)]
[167-178 (2-4)]
[169-178 (1-3)]
[178-187 (1-2)]
[178-188 (1-2)]
[178-189 (1)]
[179-186 (1)]
[179-187 (1)]
[179-189 (1)]
[179-199 (2-3)]
[187-196 (1-2)]
[187-198 (1-2)]
[187-199 (1-2)]
[188-198 (1-2)]
[188-199 (1-2)]
[190-196 (1)]
[190-199 (1-2)]
[191-198 (1)]
[191-199 (1-2)]
[200-208 (2)]
[213-219 (1-2)]
[230-238 (1-2)]
[238-244 (1-2)]
[241-254 (2-3)]
[259-271 (2-3)]
[260-271 (2)]
[263-269 (1)]
[264-271 (2)]
[278-293 (2-4)]
[279-293 (3-4)]
[280-293 (2-4)]
[281-293 (2-4)]
[288-293 (1-2)]
[292-301 (1-2)]
[294-303 (1)]
[294-304 (1-2)]
[294-305 (1-2)]
[294-306 (1-2)]
[305-317 (1-2)]
[306-317 (1-2)]
[307-317 (1-2)]
[307-323 (2-3)]
[318-323 (1)]
[318-327 (1-2)]
[318-328 (1-2)]
[329-335 (1)]
[336-341 (1-2)]
pH 3.0
pH 3.5
pH 4.0
pH 4.5
pH 6.5
%D
log time [s]

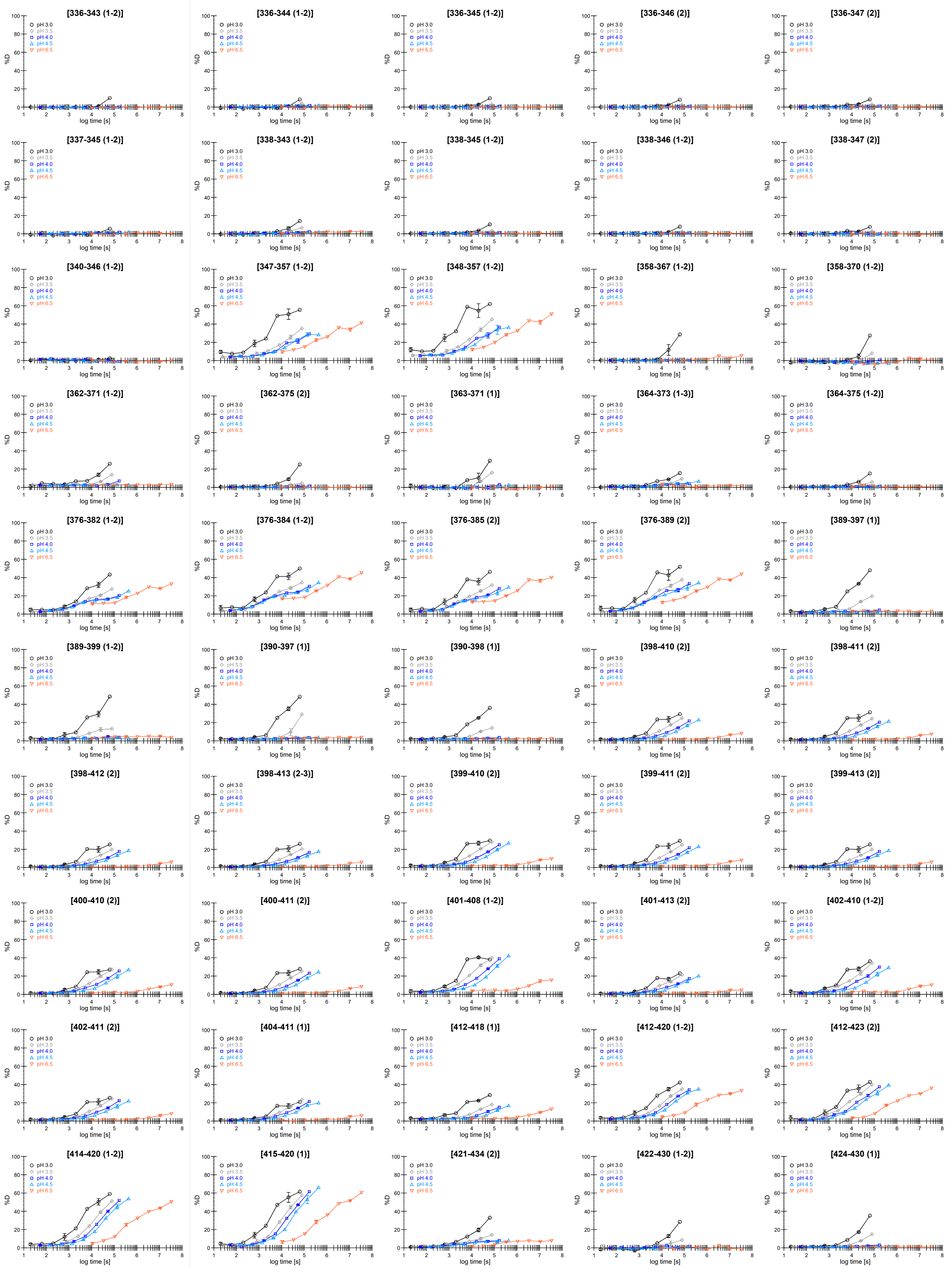
[336-343 (1-2)]
[336-344 (1-2)]
[336-345 (1-2)]
[336-346 (2)]
[336-347 (2)]
[337-345 (1-2)]
[338-343 (1-2)]
[338-345 (1-2)]
[338-346 (1-2)]
[338-347 (2)]
[340-346 (1-2)]
[347-357 (1-2)]
[348-357 (1-2)]
[358-367 (1-2)]
[358-370 (1-2)]
[362-371 (1-2)]
[362-375 (2)]
[363-371 (1)]
[364-373 (1-3)]
[364-375 (1-2)]
[376-382 (1-2)]
[376-384 (1-2)]
[376-385 (2)]
[376-389 (2)]
[389-397 (1)]
[389-399 (1-2)]
[390-397 (1)]
[390-398 (1)]
[398-410 (2)]
[398-411 (2)]
[398-412 (2)]
[398-413 (2-3)]
[399-410 (2)]
[399-411 (2)]
[399-413 (2)]
[400-410 (2)]
[400-411 (2)]
[401-408 (1-2)]
[401-413 (2)]
[402-410 (1-2)]
[402-411 (2)]
[404-411 (1)]
[412-418 (1)]
[412-420 (1-2)]
[412-423 (2)]
[414-420 (1-2)]
[415-420 (1)]
[421-434 (2)]
[422-430 (1-2)]
[424-430 (1)]
pH 3.0
pH 3.5
pH 4.0
pH 4.5
pH 6.5
%D
log time [s]

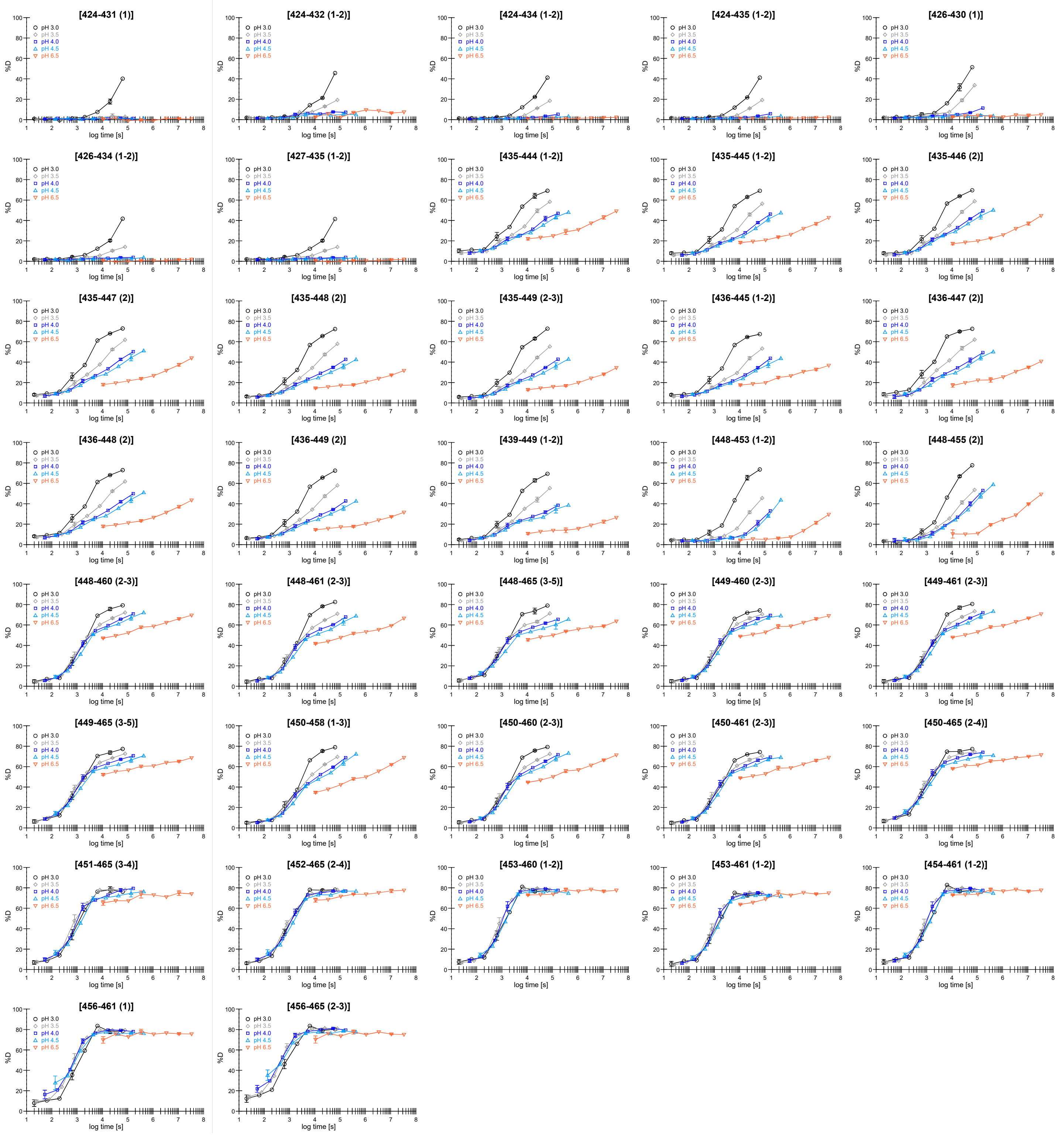
[424-431 (1)]
[424-432 (1-2)]
[424-434 (1-2)]
[424-435 (1-2)]
[426-430 (1)]
[426-434 (1-2)]
[427-435 (1-2)]
[435-444 (1-2)]
[435-445 (1-2)]
[435-446 (2)]
[435-447 (2)]
[435-448 (2)]
[435-449 (2-3)]
[436-445 (1-2)]
[436-447 (2)]
[436-448 (2)]
[436-449 (2)]
[439-449 (1-2)]
[448-453 (1-2)]
[448-455 (2)]
[448-460 (2-3)]
[448-461 (2-3)]
[448-465 (3-5)]
[449-460 (2-3)]
[449-461 (2-3)]
[449-465 (3-5)]
[450-458 (1-3)]
[450-460 (2-3)]
[450-461 (2-3)]
[450-465 (2-4)]
[451-465 (3-4)]
[452-465 (2-4)]
[453-460 (1-2)]
[453-461 (1-2)]
[454-461 (1-2)]
[456-461 (1)]
[456-465 (2-3)]
pH 3.0
pH 3.5
pH 4.0
pH 4.5
pH 6.5
%D
log time [s]

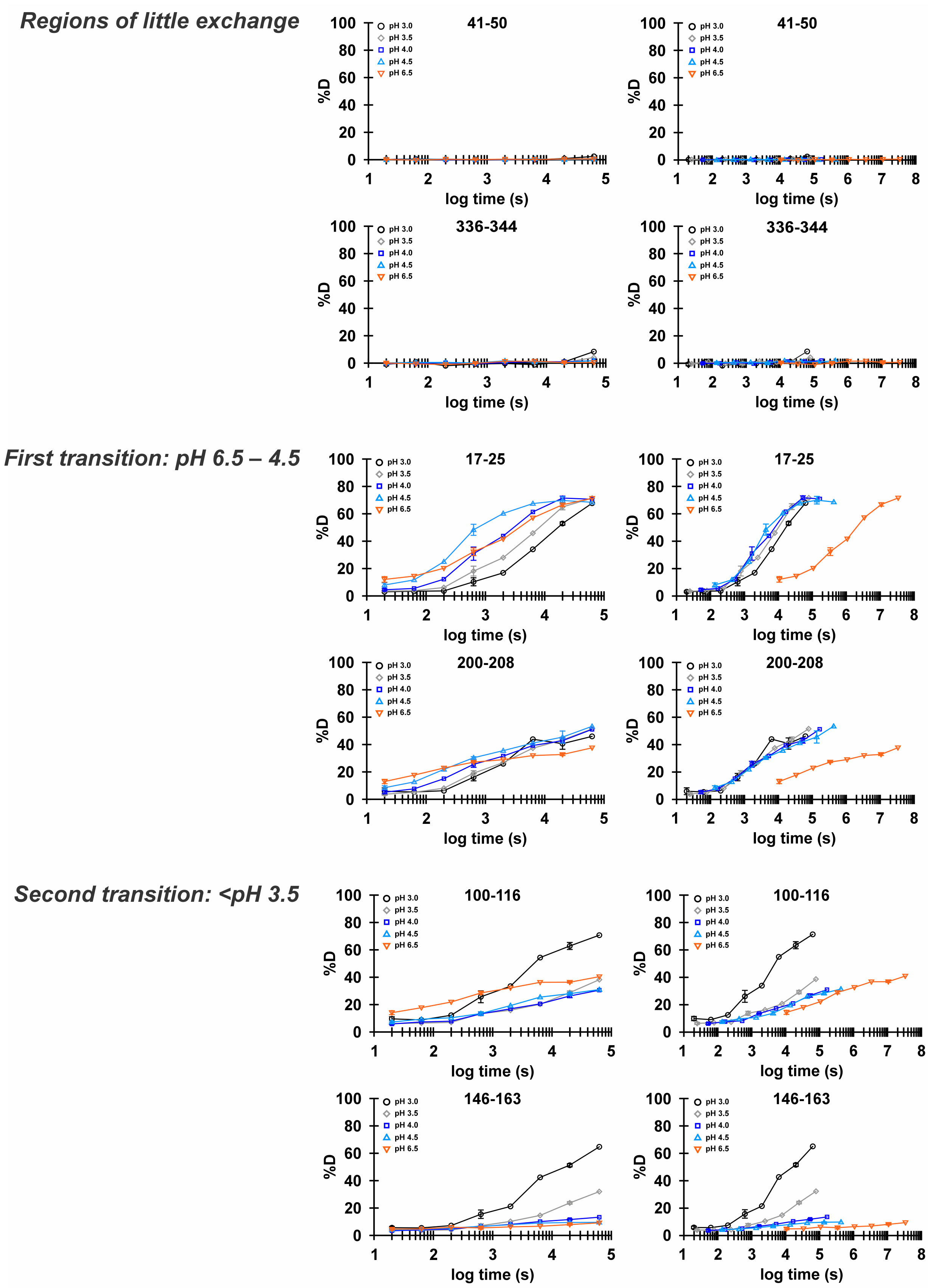

**Supplementary Figure 3C**: **Complete set of deuterium uptake curves from the HDX-MS experiment – Part C**
Selected deuterium uptake plots for the CLC-ec1 peptides shown in Figure 2c. Comparison of data without pH correction (*left panels*) and with correction factors calculated using the SPHERE server (*right panels*). Experimental conditions: pH 3.0 (black), pH 3.5 (grey), pH 4.0 (blue), pH 4.5 (cyan), and pH 6.5 (orange).

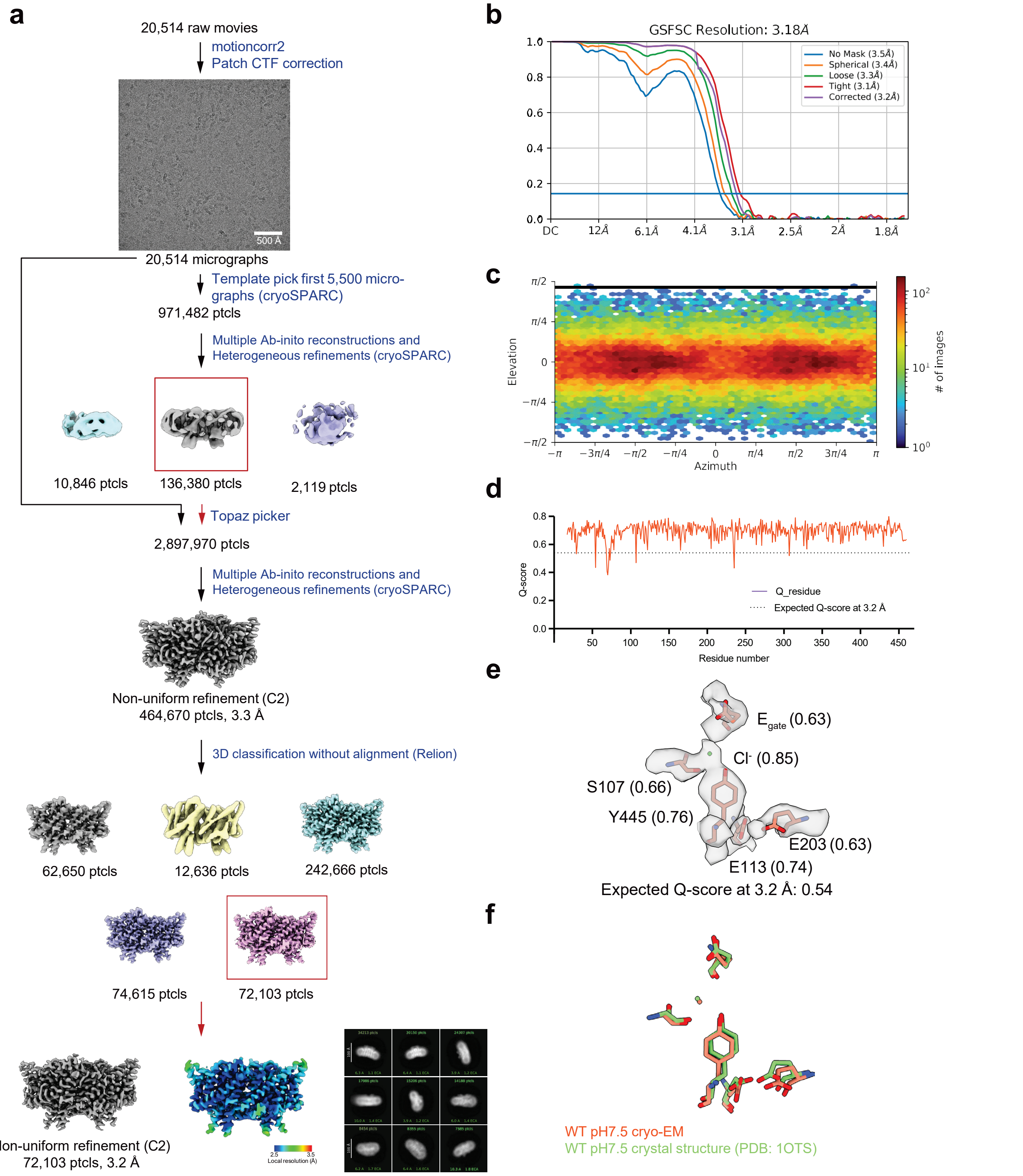

**Supplementary Figure 4: Cryo-EM workflow and validation data for CLC-ec1 at pH 7.5**. (**a**) Cryo-EM data processing workflow. The final non-uniform refinement, the local resolution estimation, and the 2D classes from the final particles are shown. (**b**) Gold-standard FSC curve. The resolution is estimated based on FSC at 0.143. (**c**) Angular distribution plot (**d**) Per-residue Q-score as a function of residue number. The expected Q-score at the map resolution is indicated by the dotted line. (**e**) The cryo-EM density and molecular model overlay for central $Cl^-$ binding site. The key residues and bound $Cl^-$ and their corresponding Q-score are annotated. (**f**) The overlay of our cryo-EM structure (salmon color) with the previous crystal structure (green) (PDB: 1OTS).

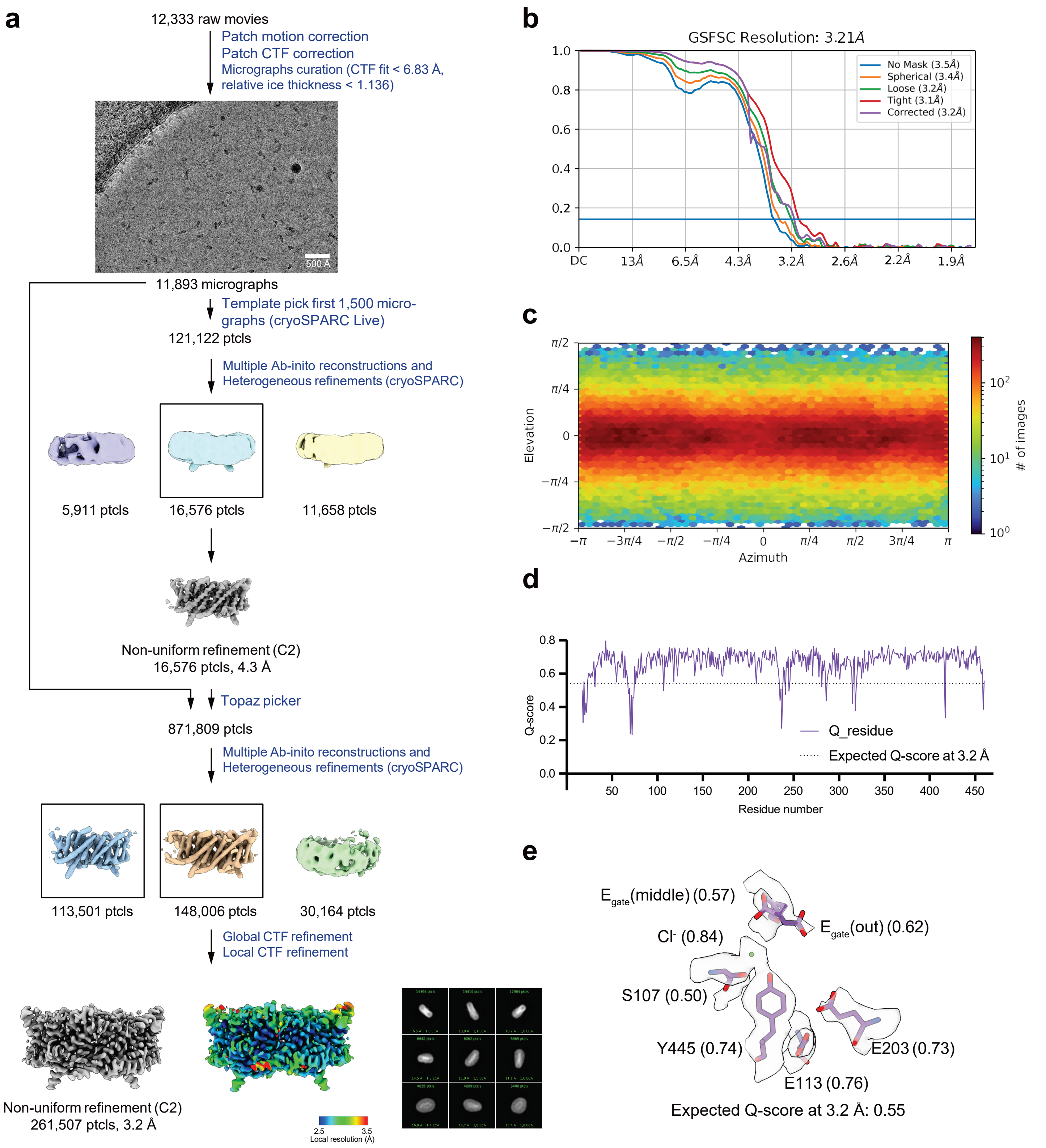

**Supplementary Figure 5: Cryo-EM workflow and validation data for CLC-ec1 at pH 4.0**. (**a**) Cryo-EM data processing workflow. The final non-uniform refinement, the local resolution estimation, and the 2D classes from the final particles are shown.(**b**) Gold-standard FSC curve. The resolution is estimated based on FSC at 0.143. (**c**) Angular distribution plot (**d**) Per-residue Q-score as a function of residue number. The expected Q-score at the map resolution is indicated by the dotted line. (**e**) The cryo-EM density and molecular model overlay for central $Cl^-$ binding site. The key residues and bound $Cl^-$ and their corresponding Q-score are annotated.

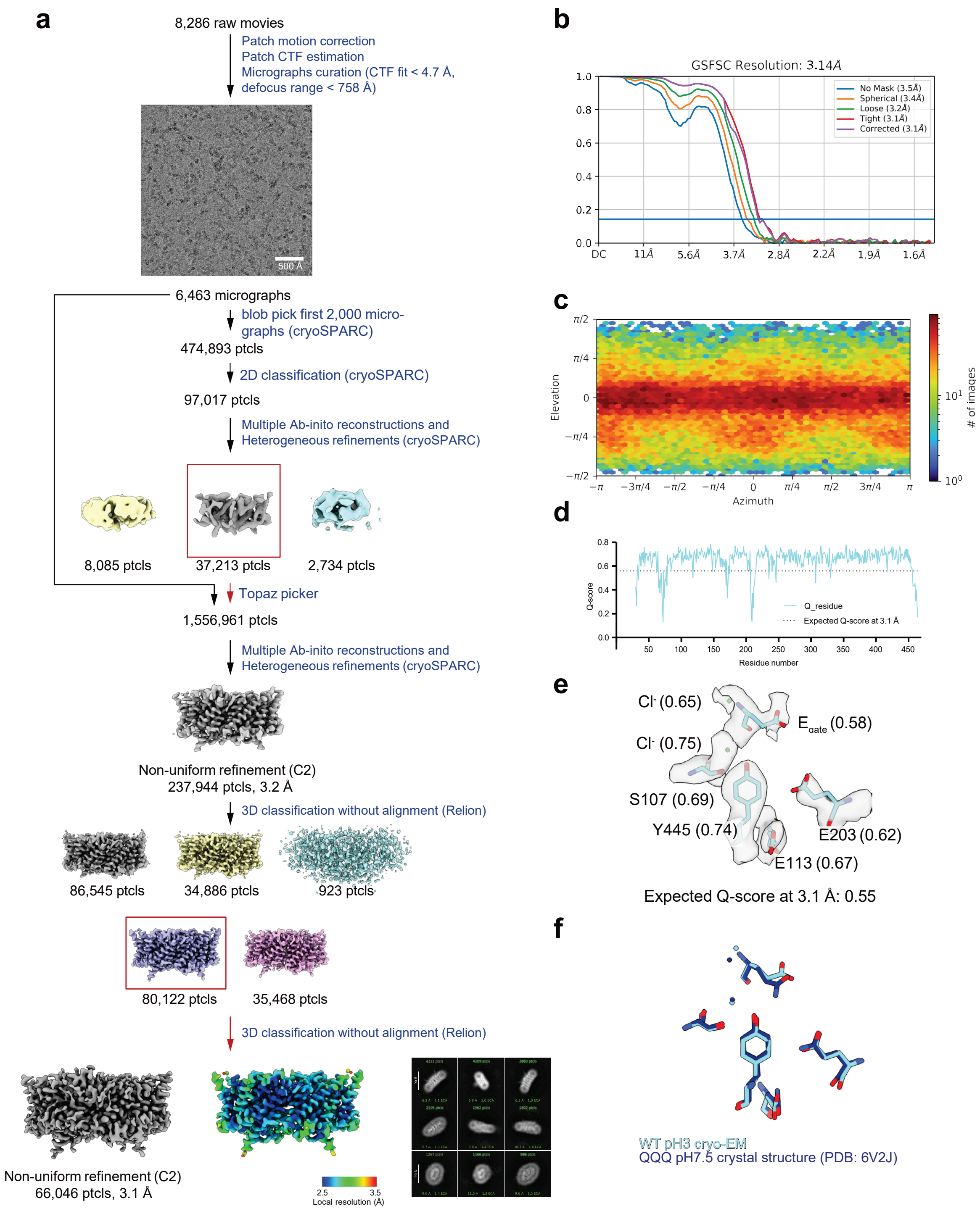

**Supplementary Figure 6: Cryo-EM workflow and validation data for CLC-ec1 at pH 3.0**. (**a**) Cryo-EM data processing workflow. The final non-uniform refinement, the local resolution estimation, and the 2D classes from the final particles are shown. The ~160,000 particles that were discarded during 3D classification in Relion could be reconstructed to comparable nominal resolution as our final model; however, the resulting maps showed missing or incomplete density in several regions of the protein. Specifically, density was missing for the I–J loop, the B–C loop, and helix D. To enable building a complete atomic model, we focused on the subset of 80,122 particles that yielded the most complete map. The absence of density in the excluded classes reflects increased flexibility in these regions, consistent with our HDX results that conformational dynamics are enhanced at pH 3.0. (**b**) Gold-standard FSC curve. The resolution is estimated based on FSC at 0.143. (**c**) Angular distribution plot (**d**) Per-residue Q-score as a function of residue number. The expected Q-score at the map resolution is indicated by the dotted line. (**e**) The cryo-EM density and molecular model overlay for central $Cl^-$ binding site. Key residues and bound $Cl^-$ and their corresponding Q-score are annotated. (**f**) Overlay of our cryo-EM structure (cyan) with the previous QQQ crystal structure (dark blue) (PDB: 6V2J).

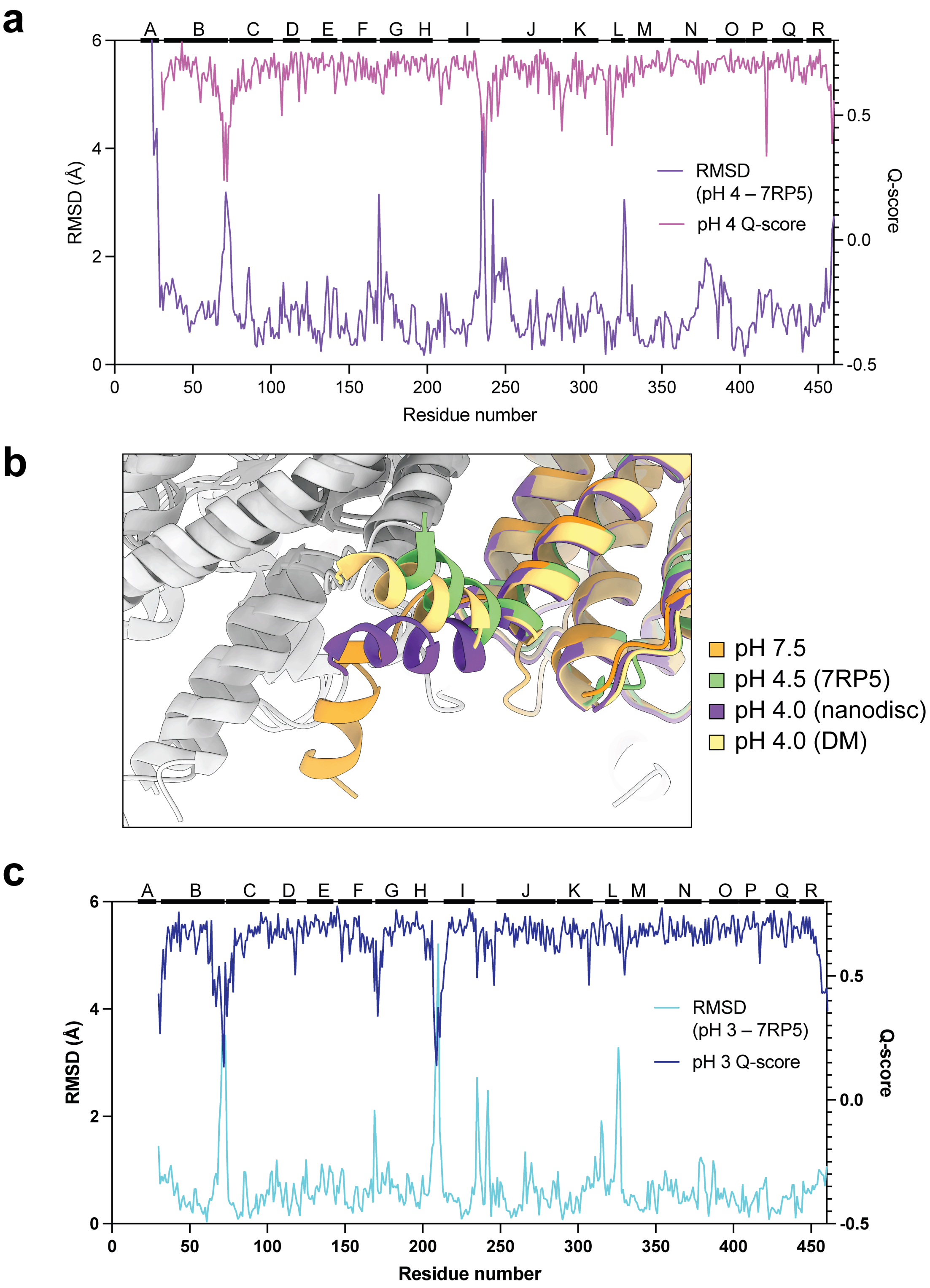

**Supplementary Figure 7: Comparison to pH 4.5 structure, PDBID: 7RP5.** (**a**) RMSD between 7RP5 and the pH 4.0 structure presented here. The large changes are predominantly in loop regions where Q-scores are low and therefore model-to-model comparisons have low certainty (**b**) Structure overlay showing change in orientation of helix A in the pH 4.0 structure compared to pH 7.5 and to pH 4.5 (7RP5). Our additional structure at pH 4.0 in DM shows that the orientation of helix A depends on the membrane mimetic used. (**c**) RMSD between the pH 3.0 structure and 7RP5 (pH 4.5).

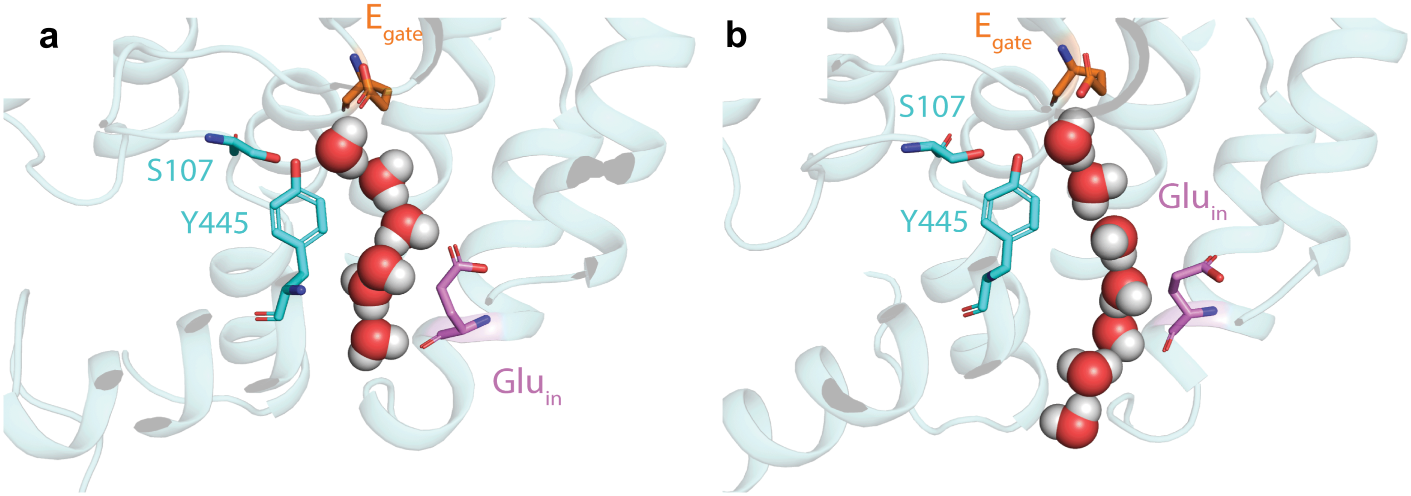

**Supplementary Figure 8: Examples of water wires with (a) and without (b) connection to $Glu_{in}$.**

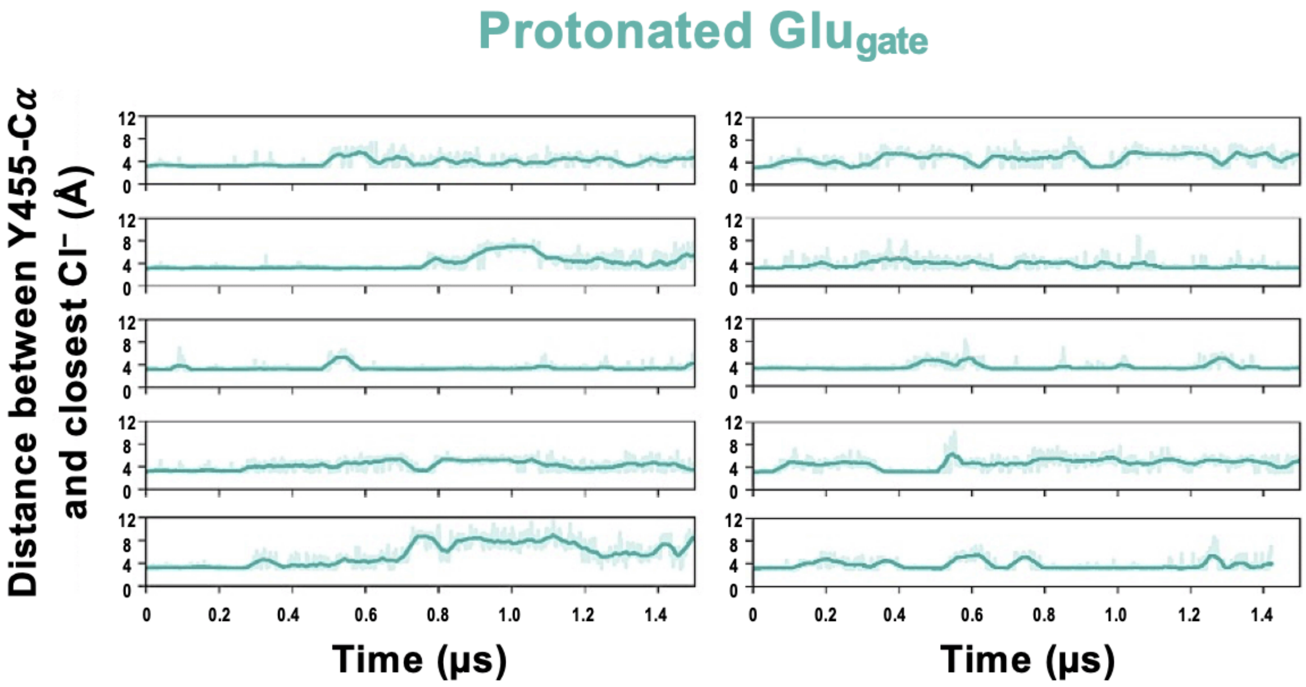

**Supplementary Figure 9: $Cl^-$ remains in the anion pathway during simulations with $E_{gate}$ protonated.** The presence of $Cl^-$ within the anion pathway during simulations was evaluated by plotting the distance from the Cα atom of inner-gate residue Y445 to the nearest $Cl^-$ ion.

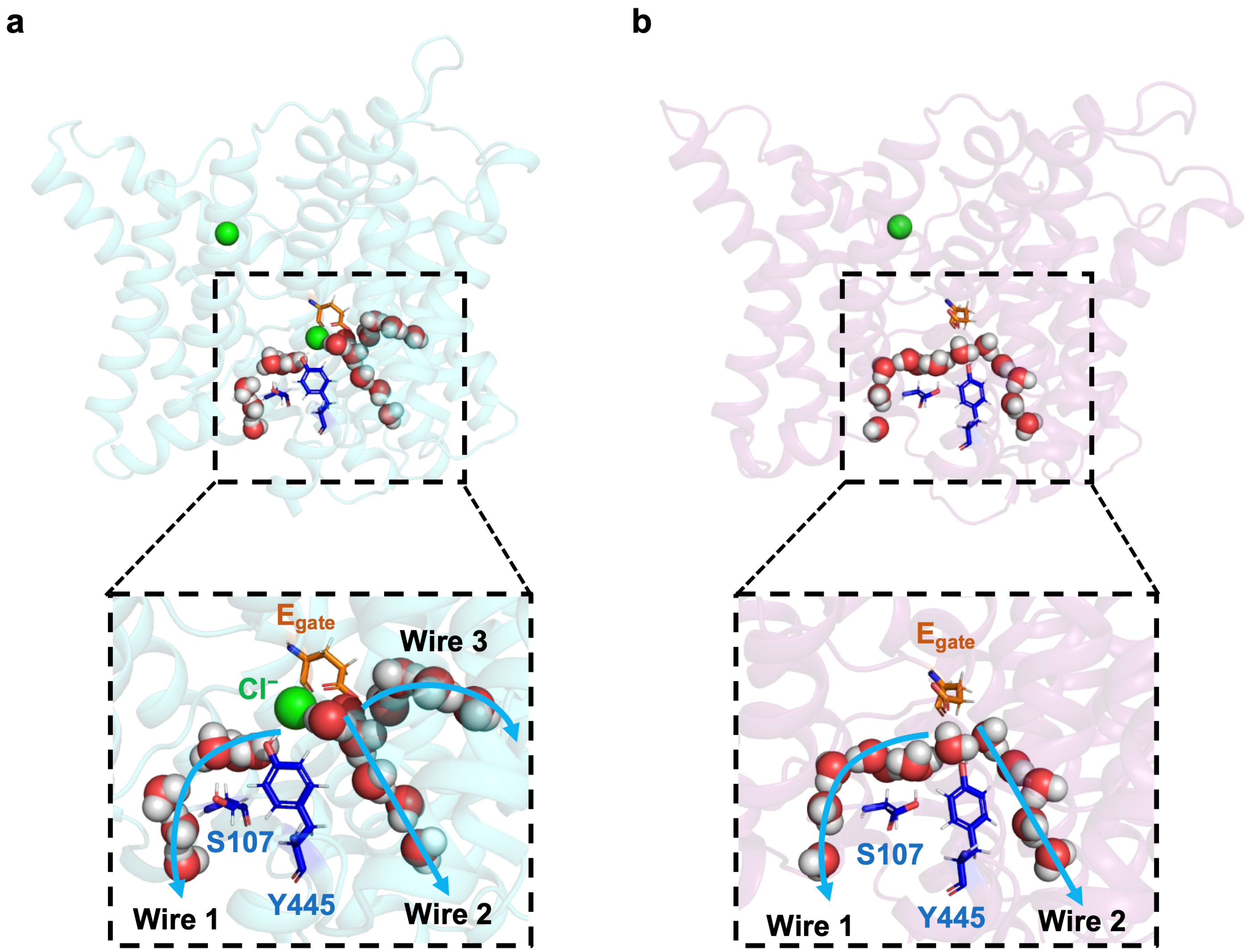

**Supplementary Figure 10: Water wires occur in the presence and absence of Cl⁻.** (**a**) Example snapshot of water wires formed in the presence of Cl⁻ in the permeation pathway. (**b**) Example snapshot of water wires formed in the absence of Cl⁻ in the permeation pathway.

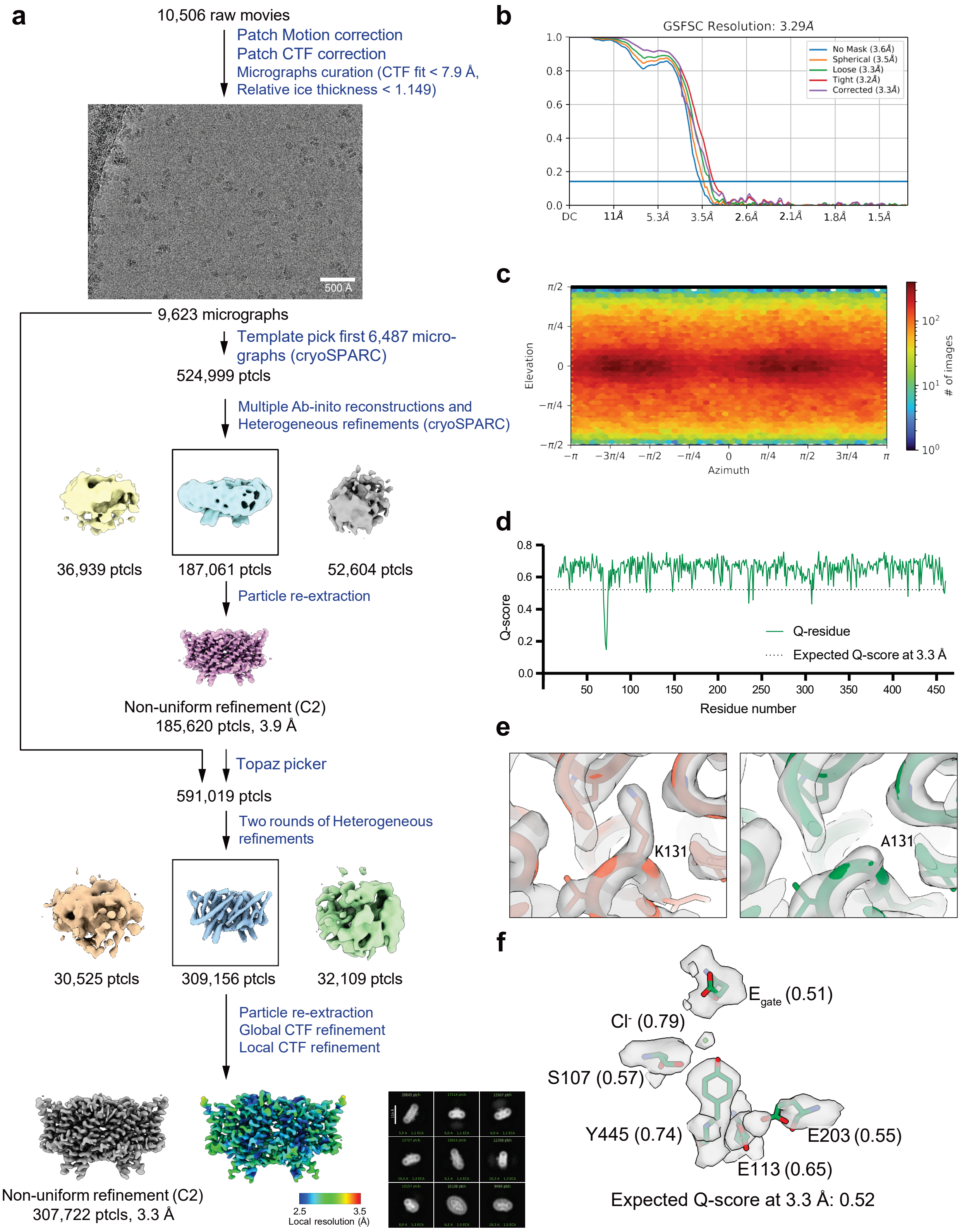

**Supplementary Figure 11: Cryo-EM validation data for K131A CLC-ec1.** (**a**) Cryo-EM data processing workflow. The final non-uniform refinement, the local resolution estimation, and the 2D classes from the final particles are shown. (**b**) Gold-standard FSC curve. The resolution is estimated based on FSC at 0.143. (**c**) Angular distribution plot (**d**) Per-residue Q-score as a function of residue number. The expected Q-score at the map resolution is indicated by the dotted line. (**e**) The cryo-EM density and molecular model overlay in the region of K131 (left panel) and A131 (right panel). (**f**) The cryo-EM density and molecular model overlay for central $Cl^-$ binding site. The key residues and bound $Cl^-$ and their corresponding Q-score are annotated.

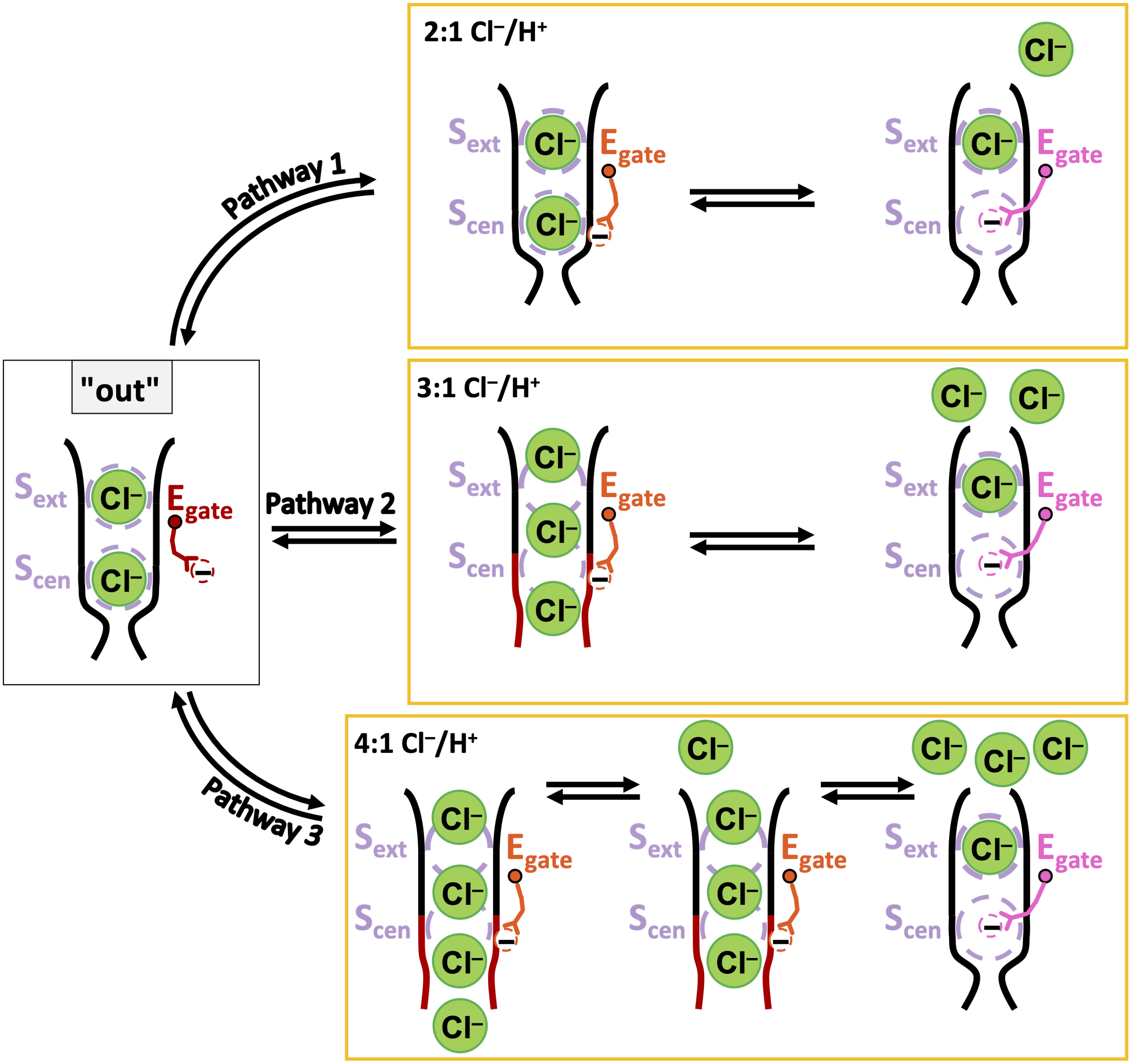

**Supplementary Figure 12: Uncoupling in K131 mutant CLC-ec1 transporters.** The mild uncoupling that occurs in K131 mutants, with $Cl^-/H^+$ stoichiometry increased from 2 to ~3, could result from a moderate destabilization of the inner gate. In the normal 2:1 transport cycle (pathway 1, model at top), the inner gate opens only after Egate has fully entered the permeation pathway. In K131 mutants, the slightly destabilized inner gate opens prematurely, permitting 1 or 2 Cl- to slip through before Egate blocks the pathway (pathways 2 and 3). The observed 3:1 $Cl^-/H^+$ stoichiometry can be produced solely via pathway 2 or by a mixture of pathways that averages to 3:1.

**Supplementary Information References**

1. Bai Y, Milne JS, Mayne L, Englander SW. Primary structure effects on peptide group hydrogen exchange. *Proteins* **17**, 75-86 (1993).

2. Connelly GP, Bai Y, Jeng MF, Englander SW. Isotope effects in peptide group hydrogen exchange. *Proteins* **17**, 87-92 (1993).

3. Moroco JA*, et al.* High-Throughput Determination of Exchange Rates of Unmodified and PTM-Containing Peptides Using HX-MS. *Mol Cell Proteomics* **24**, 100904 (2025).
